# Infer global, predict local: quantity-quality trade-off in protein fitness predictions from sequence data

**DOI:** 10.1101/2022.12.12.520004

**Authors:** Lorenzo Posani, Francesca Rizzato, Rémi Monasson, Simona Cocco

**Author notes:** Co-senior authors (Rémi Monasson, Simona Cocco). To whom correspondence should be addressed. (SC), (LP).

## Abstract

Predicting the effects of mutations on protein function is an important issue in evolutionary biology and biomedical applications. Computational approaches, ranging from graphical models to deep-learning architectures, can capture the statistical properties of sequence data and predict the outcome of high-throughput mutagenesis experiments probing the fitness landscape around some wild-type protein. However, how the complexity of the models and the characteristics of the data combine to determine the predictive performance remains unclear. Here, based on a theoretical analysis of the prediction error, we propose descriptors of the sequence data, characterizing their quantity and quality relative to the model. Our theoretical framework identifies a trade-off between these two quantities, and determines the optimal subset of data for the prediction task, showing that simple models can outperform complex ones when inferred from adequately-selected sequences. We also show how repeated subsampling of the sequence data allows for assessing how much epistasis in the fitness landscape is not captured by the computational model. Our approach is illustrated on several protein families, as well as on in silico solvable protein models.

**Significance Statement:** Is more data always better? Or should one prefer fewer data, but of higher quality? Here, we investigate this question in the context of the prediction of fitness effects resulting from mutations to a wild-type protein. We show, based on theory and data analysis, that simple models trained on a small subset of carefully chosen sequence data can perform better than complex ones trained on all available data. Furthermore, we explain how comparing the simple local models obtained with different subsets of training data reveals how much of the epistatic interactions shaping the fitness landscape are left unmodeled.

## Introduction

Predictability of evolution of organisms in fitness landscape has been a driving concept in the development of evolutionary biology since the origins of the field (1–5). In particular, our capability to predict the effects of detrimental mutations has enormous practical impact on the diagnosis of genetic variances causing diseases (6–10). This issue can now be quantitatively investigated, thanks to high-throughput sequencing and mutagenesis experiments, which allow for in-vivo and in-vitro measurements of the effects of many mutants (1, 5, 11–25). However, despite the impressive progress of these large-scale techniques, the number of possible mutations, growing exponentially with the protein length, is so huge that measuring the fitness landscape in its entirety is out of reach, with the exception of short protein regions (1). Computational approaches, in particular machine-learning-based models exploiting the large corpus of available sequence data (26, 27) are needed for the full reconstruction and prediction of fitness landscapes. Briefly speaking, these methods are based on the assumption that statistically rare mutations (in homologous sequence data) are likely to be deleterious (6, 28). Such conservation-based methods can be combined with structural (7, 29), physico-chemical (8), as well as phylogenetic (30, 31) information.

Graphical Potts models, also called direct coupling analysis (DCA)(32–34), have pushed further the approaches based on sequence conservation by including statistical couplings capturing pairwise amino-acid covariation. These couplings allows DCA to account for background effects on the mutations depending on the wild-type (*wt*) sequence under consideration. DCA is thought to approximate the fitness landscape reflecting the structural and functional properties common to homologous proteins. As sketched in Fig. 1, natural sequences are assumed to lie at, or close to the different peaks of the fitness landscape explored during evolution. The scores of sequences around the *wt* protein provide predictions for the effects of single or multiple mutations on fitness, in good agreement with mutational effects measured through mutagenesis experiments (34–38). Other approaches for fitness prediction exploit deep learning (DL) architectures, at the origin of recent progress in image or natural language processing, as well as in protein folding (9, 10, 39–41). DL models have much higher expressive power than pairwise graphical models, but demand massive sequence data to be trained. Recent applications of DL to protein fitness modelling combine unsupervised learning of hundreds of millions of sequences with supervised learning of mutagenesis experimental data (38, 42, 43).

**Fig. 1.**
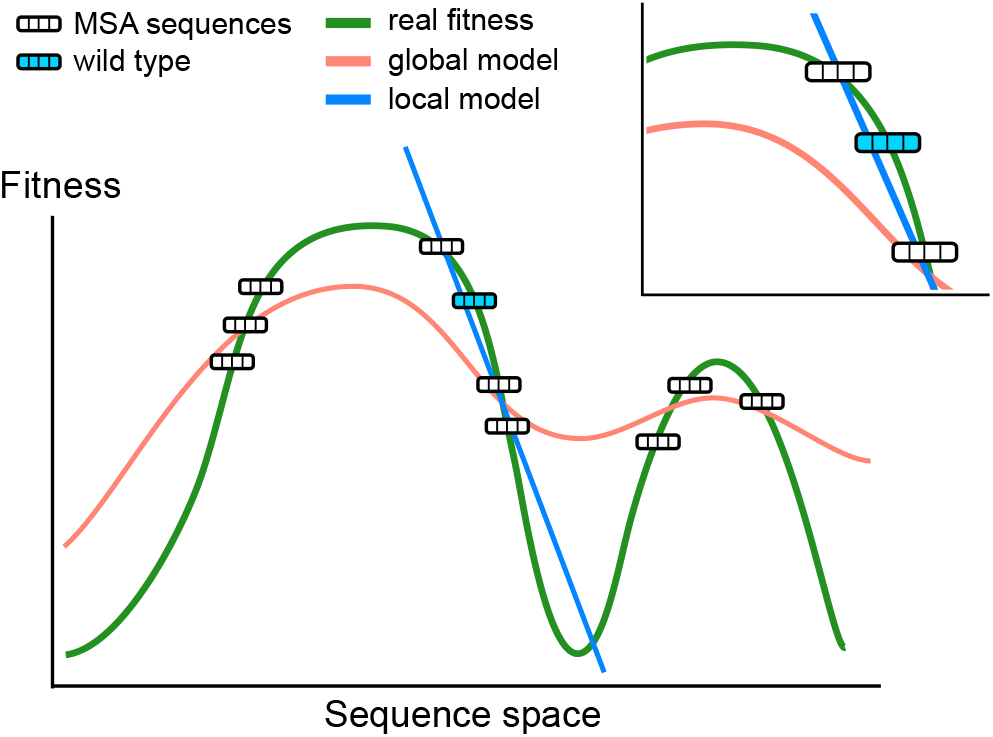
Schematic visualization of the fitness landscape over the sequence space (grey dashed line). Two models (red and blue curves) are inferred to assign high fitness values to sequences found in the Multi-Sequence Alignment (MSA) of a protein family. A complex model (red curve) can be a better predictor of the landscape globally, while scoring poorly in predicting single-point mutations around a specific wild-type sequence, see local fitness landscape in the zoomed area. Conversely, a simple model (blue line) fitted on a local subset of sequences can give a better local approximation of the landscape, but will likely fail in distant regions of the fitness landscape.

Depending on the protein family under consideration, multisequence alignments (MSA) show huge variations in sizes, with tens to hundred of thousands sequences, and in homology, ranging from ~ 30%, for alignments of orthologous sequences (27, 37, 44), to 90%, for HIV sequences of the same clade (35, 45). The quantity and quality (diversity) of the data, as well as the models considered are empirically known to strongly impact the performances for fitness prediction. As pointed out in (46), classical methods based on homology detection, such as SIFT (6), PolyPhen-2 (7), Align-GVGD (8), rely on different empirical procedures in selecting the alignments, and are not always optimal. Remarkably, single mutations effects are predicted with comparable accuracy by graphical models inferred from a small number of highly similar sequences of the HIV envelope protein (35) and from a much larger number of diverse sequences of Betalactamases, while the two proteins have comparable lengths (37). Gemme, a recently introduced algorithm based only on conservation and phylogenetic tracing of mutations (31) was shown to outperform deep neural networks models (39) in predicting the effect of mutations in viral sequences, all characterized by a large degree of similarity. Furthermore, it is known that the performance of models trained from Uniprot sequences with high pairwise alignment score to a fixed *wt* sequence considerably vary with the threshold used for alignment (37, 46).

These examples suggest the existence of a compromise between taking into account many sequence data to get statistics and removing far away sequences, whose relation to fitness may be very different from *wt* due to complex epistatic effects. This compromise, in turn, depends on the expression power of the model considered, which can be tuned at will, and on the complexity of the fitness landscape, which is generally unknown. As sketched in Fig. 1, on the one hand, predicting mutations around the *wt* requires *local* reconstruction of the landscape only, a task within reach of simple models with few defining parameters. These models are however unreliable for sequences far away from the *wt* sequence; hence, only few data points, concentrated around the latter can be actually used for training. On the other hand, powerful models able to capture the complex features, such as high-order epistasis that characterize the *global* fitness landscape on large scales can, in principle, exploit at best sequence data. However, even if the available data are sufficient to infer their huge number of parameters with enough accuracy, it is unclear whether the global description they offer allows for an accurate local reconstruction of the fitness landscape around the *wt* protein. The scope of the present work is to provide theoretical foundation to address this question.

Careful analysis of the different contributions to the prediction error allows us to quantitatively understand how fitness prediction performance depend on both model complexity and on the quality/quantity of data, and to estimate the amount of ‘complexity’ in the fitness landscape that is not captured by the model. Our theory is in full agreement with the analysis of sequence data and mutagenesis experiments for 7 protein families we have studied. We also validate our approach *in silico* on Lattice-Protein models (47–50), for which the groundtruth for the fitness is mathematically well defined. Last of all, we demonstrate how our framework allows us, in practice, to optimally tune sequence alignments and models to maximize the performance in fitness prediction.

## Results

### A. Quantity-quality trade-off in MSA sequence selection

We consider a reference sequence, hereafter referred to as *wt*. We denote by 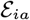 the variation of fitness resulting from the mutation *wt_i_* → *a* on the *i^th^* site of *wt*. This quantity can be estimated experimentally, either in vivo (relative enrichment of organisms with mutated gene compared to *wt*), or in vitro (measurement of appropriate biochemical property).

A computational model provides a predictor, 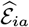, for the difference of fitness between the mutant and the *wt*. The overall quality of the predictor will be assessed through the Spearman coefficient *ρ* between the mutation effects computed with the model 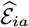 and with the experimental data 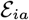. Using Spearman correlations allows one to capture monotonous relations, irrespective of non-linearities.

The computational model is generally trained from homologous sequences to *wt*, *i.e*. belonging to the same protein family. The similarities between the *wt* and these sequences, sampled from evolutionary diverse organisms, can vary significantly. As an illustration, we consider the RNA binding domain of the nuclear poly(A)-binding protein (PABPN1), involved in the synthesis of the mRNA poly(A) tails in eukaryotes (14). Any two sequences in the corresponding MSA (as used in (37)) generally have few amino acids in common (mean Hamming distance -normalized by sequence length- between pairs of sequences in the MSA = 0.75). As a result, a specific sequence, such as the *wt* of *Saccharomyces cerevisiae*, is generally surrounded by a small number of similar sequences and is far away from most of the MSA (RNA-bind protein: mean normalized Hamming distance between *wt* and MSA sequences = 0.73, Fig. 2A; see Supplementary Fig. 3 for similar results on other families).

**Fig. 2.**
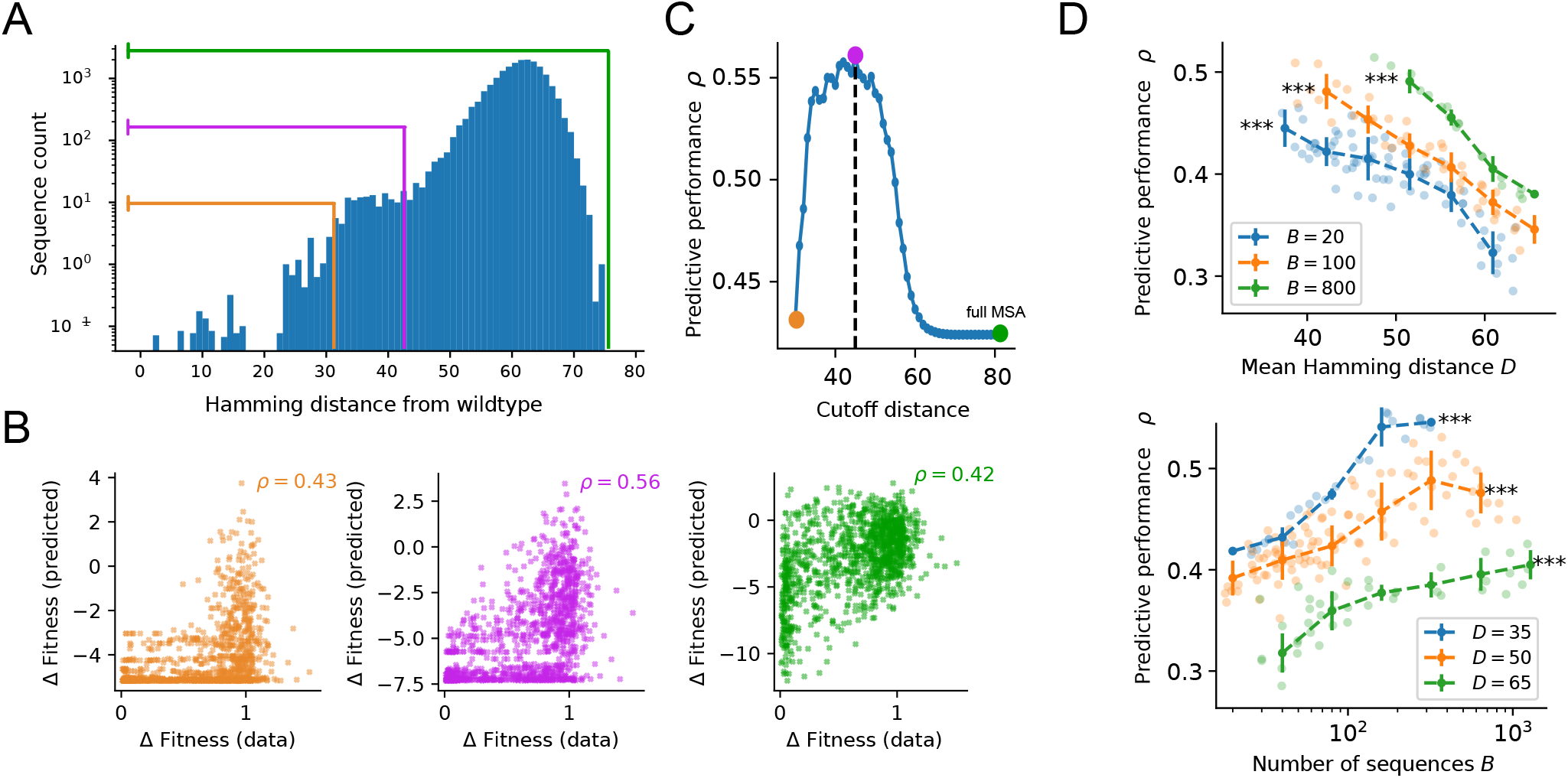
Behaviour of model predictive performance with different selections of training data. **A.** Distribution of Hamming distances to the *wt* sequence (RNA-binding domain of Pab1-Yeast) in the MSA of (37). Note the log scale on the *y* axis. The three colored lines correspond to three possible sequence selections performed by excluding sequences farther than a certain threshold *d_cut_* from *wt*. A smaller *d_cut_* corresponds to fewer sequences with a lower mean Hamming distance to the *wt*, denoted as *D*. **B** Comparison between predicted and experimental fitness mutational effects for an independent-site model trained on the three sub-MSAs corresponding to, respectively, *d_cut_* = 32 (orange), 43 (purple), and 82 (green). The Spearman correlation coefficient *ρ* between predicted and experimental values defines the predictive performance of the model. **C** Same analysis as panel B repeated for all possible cutoffs between *d_cut_* = 32 and *d_cut_* = 82 (the sequence length). The non monotonous behavior of the predictive performance indicates that a trade-off between number of sequences (denoted as *B*) and proximity to *wt* is controlling the predictive performance of the inferred model. **D.** Systematic analysis of the predictive power *ρ* as a function of the mean Hamming distance *D* of sub-alignments with fixed size *B* (top), and of the sub-alignment size *B* at fixed Hamming distance *D* (bottom). Each individual point shows the average over *n* = 5 sub-samples obtained at the corresponding values of *D* and *B* (see Methods). The dashed curves and error bars are computed by binned average and standard deviation over the displayed individual points. All significance levels refer to Spearman rank correlation of the individual points. *** P<0.001.

Hereafter, we show that sequences far away from *wt* are of poor quality for fitness prediction. To do so we train independent-site Potts models (Methods) on shorter MSAs obtained by discarding sequences further than a certain cut-off distance *d_cut_* from *wt*. As *d_cut_* becomes smaller, fewer sequences with higher proximity are selected (Fig. 2A). We see that the performance consistently increases when decreasing the cut-off distance, up to a peak *ρ* = 0.56 at *d_cut_* = 43, a 33% increase with respect to the full MSA (*ρ*(*d_cut_* = 82) = 0.42), see Fig. 2B,C). After peaking, the performance starts decreasing again due to the increasingly-lower number of sequences in the MSA, see Fig. 2C.

The non monotonous behavior of the predictive performance indicates that a trade-off between the number of sequences and their proximity to *wt* is controlling the predictive performance of the inferred model. To investigate the respective effects of these two quantities, we create sub-alignments of the original MSA with controlled sizes *B* (effective number of sequence taking into account sequence redundancy, see (51) and Methods) and average Hamming distances to *wt*, which we denote as *D*. We then test how the performance of the independent-site Potts model trained on these sub-alignments relates to these two quantities.

This analysis showed that the predictive performance strongly depends on the mean Hamming distance *D* and on the number *B* of sequences (P<0.001 for all Spearman correlations between *ρ* and *B* or *D*, Fig. 2D). The performance significantly decreases with *D* at fixed *B*, *i.e*. when the quality of the data deteriorates and their quantity is kept fixed, and increases with *B* at fixed *D*, *i.e*. when the quantity of data increases at fixed quality. Similar results are found for the six other protein families under study (see SI Fig. 1).

### B. Theoretical investigation of the quantity-quality trade-off

To study the trade-off between quality and quantity we draw our inspiration from the bias-variance framework developed in statistics (52, 53). Let us consider the error 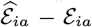 between the statistical predictor 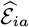 and the experimental fitness 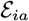. This error can be decomposed into the sum of two contributions: (1) a systematic bias in the prediction, due to the inability of the model to capture the exact relation between sequence mutation and fitness, (2) a statistical error coming from the fact that the predictive model has been trained on a particular data set; the value of this contribution fluctuates when the data set changes, and is expected to be smaller and smaller for larger and larger data sets.

Consequently, the mean squared error on the single-point mutation *wt_i_* → *a* can be written as the sum of a *squared bias* and a *variance* contributions,

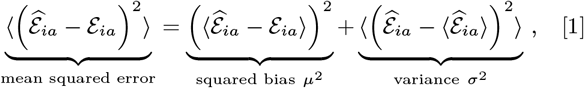

where averages 〈〉 are taken on repetitions of the prediction process in fixed conditions (quality and quantity of data). Notably, these two quantities are hard to minimize together. For instance, powerful models with many parameters will accurately fit the data and thus achieve small squared biases *μ*^2^, but will result in large variances *σ*^2^ due to the statistical errors on the many parameters to be inferred. As we will see below, we can directly relate our descriptors of quality (*D*) and quantity (*B*) of the sequence data to, respectively, the squared bias and the variance as defined in Eq. (1). Furthermore, we introduce below a class of increasingly powerful Potts models to investigate the effect of model complexity on these two quantities and, ultimately, on the predictive performance.

#### K–link Potts model

We consider hereafter the class of sparse Potts models, which include *K* pairwise couplings between the protein sites, *J_ij_*(*a, b*), whose values depend on the amino acids they carry and a field (position weight matrix) *h_i_*(*a*) on each site; These parameters are learned from the MSA (Methods). The choice of the *K* pairs of sites carrying couplings is decided based on heuristics, which aim at capturing relevant interrelations between the residues (Methods).

By tuning the value of *K*, we can interpolate between the independent-site model (*K* = 0, *i.e*. no coupling) and the full Potts model 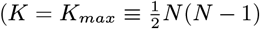 couplings, where *N* is the protein length). Imposing small values of *K* is a way to regularize the inferred network of interactions^*^.

For the *K*-link Potts model the predictor of the fitness difference resulting from the mutation *wt_i_* → *a* reads

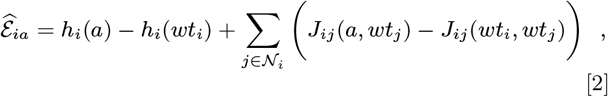

where the sums runs on the sites *j* in the neighborhood 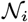of site *i, i.e*. coupled to *i* (Methods). This neighborhood is empty for the independent-site model.

#### Estimation of variance

For the Potts model, expressions for the uncertainties on the inferred fields *h_i_*(*a*) and couplings *J_ij_*(*a, b*) can be formally derived from sampling errors due to the finite size of the data set. The resulting variance 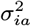 of the predictor 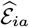 for a specific K-link model can then be estimated from Eq. (2) (38, 54), see SI Appendix A. Averaging 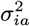 over the sites *i* and mutations *a*, we obtain a single global variance,

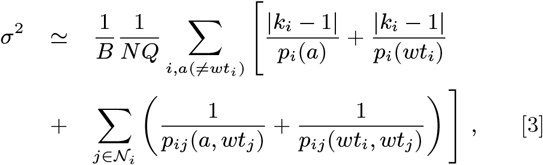

where *Q* = 20 is the number of amino-acid types, and *k_i_* is the cardinality of 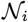, i.e. the number of sites interacting with *i* in the model. The global variance depends on the statistics of the data through the probabilities *p_i_*(*a*) of finding amino acid *a* on the *i*-th site and *p_ij_*(*a, b*) of finding simultaneously *a* on site *i* and *b* on site *j* computed on the sub-alignment. *σ*^2^ thus increases with residue conservation, due to the contributions of amino acids that are rarely observed on some sites in the sub-alignment and have low *p_i_*(*a*), and with the number *K* of coupling parameters in the model. We also see that *σ*^2^ is inversely proportional to the number of sequences, *B*. The variance therefore decreases with the *quantity* of data.

#### Estimation of squared bias

Computing the squared bias *μ*^2^ in Eq. (1) is generally hard, not to say impossible, as it requires detailed knowledge of the fitness landscape. We rely below on simplifying assumptions to gather insights on the value and meaning of the bias.

Assume first that we use the independent-site model for fitness prediction. If the ‘true’ fitness landscape shows no epistasis, this model is exact (up to statistical fluctuations due to the finite amount of training data, taken care of by *σ*^2^), and the bias vanishes. Therefore, a non zero bias would signal the presence of epistatic interactions between residues not captured by the simple model used for predictions.

This statement can be extended to more complex landscapes and models. Assume now that the fitness landscape is characterized by pairwise epistasis only, *i.e*. the fitness differences 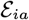 are exactly described by a full Potts model with *K_max_* interactions 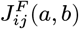 through an equation analogous to Eq. (2). The *K*–link Potts model used for fitness prediction will not be powerful enough to account for the complexity of this landscape and of the sequence data if *K* < *K_max_*. As a result a non-zero squared bias will appear, whose expression is derived in SI, Appendix B, and reads

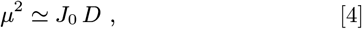

where *D* is the mean Hamming distance of the sub-alignment sequences to *wt*, and the *bias factor J*_0_ is the product of a multiplicative factor depending on the background distribution of amino acids in the MSA and of the variance of the epistatic couplings *J^F^ not included in the prediction model. J*_0_ is thus a decreasing function of *K*.

This expression of *μ*^2^ confirms that the Hamming distance *D* is related to the notion of *quality* (relative to the *wt*) of the sequence data, as varying *D* affects the systematic error (bias) of the predictive model.

### C. Validation of the theory on Lattice Proteins

To validate the key role of the squared bias and of the variance in explaining performance, as well as their approximate expressions above and the interpretation of the bias factor *J*_0_ as reflecting unmodelled epistasis, we resort to an in silico model for proteins folding on a 27-site cubic lattice (47, 49, 50, 55, 56), see Fig. 3A. In the model, the fitness represents the propensity of a protein sequence to fold into one specific conformation, called native, out of the ≃ 10^5^ folds on the cube (49). As we can precisely compute the exact value of the fitness, the ground-truth values of the squared bias and of the variance defined in Eq. (1) can be computed with great accuracy (see Methods); we hereafter denote these ground truth values by 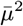 and 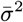.

**Fig. 3.**
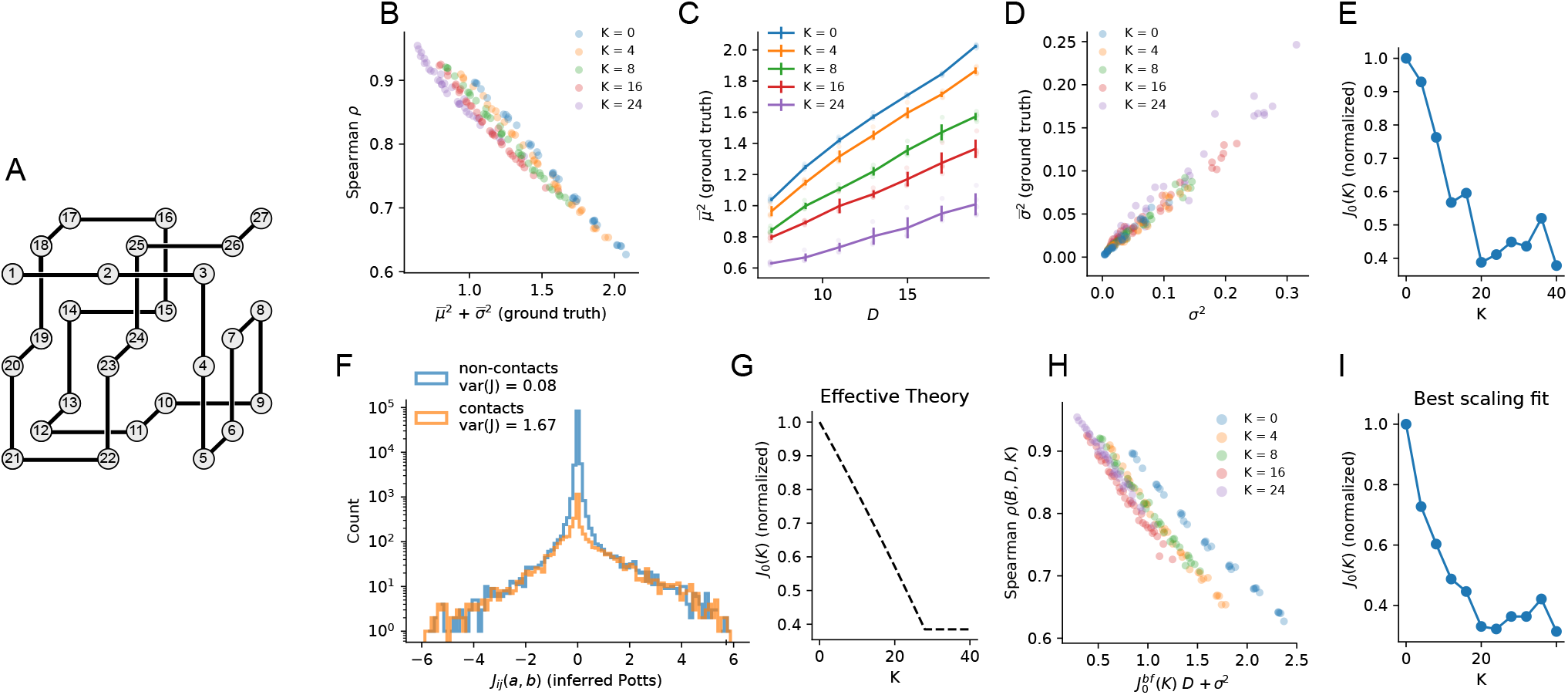
Quantity-quality trade-off for lattice proteins. **A**: Cubic fold that defines the protein family in the lattice model. Amino acids on sites that are in proximity to each other interact, and define the energy of the protein (Methods). **B**: Predictive performance *ρ* for single mutations of 5 Sparse Potts models with different degrees of sparsity (defined by *K*, the number of pairwise links included in the energy function; K=0 is the independent model) vs. 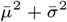. The collapse of the results is in agreement with Eq. (5). **C**: Squared bias 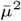 vs. mean Hamming distance in the sequence data, see Eq. (4), for the same sparse Potts models as in panel B. Line plots and error bars show mean and standard deviation at a given *D* and different *B*s. **D**: Variance 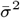 vs. estimated variance *σ*^2^ in Eq. (3) for the same Sparse Potts models as in panel B. **E**: Bias factor *J*_0_(*K*) (divided by *J*_0_(0)) obtained by fitting the squared bias as a linear function of the mean Hamming distance for the various K-link models in panel C. **F**: Visualization of pairwise couplings inferred by a fully-connected Potts model, highlighting the larger variance of couplings associated to structural contacts (in orange) compared to non-structural ones (in blue) - note the log scale on the y axis. **G**: Normalized value of *J*_0_ (*K*) (divided by *J*_0_ (0)) obtained with an effective theory using the variance of couplings associated to modeled and un-modeled structural contacts, see Appendix A. **H**: scaling for predictive performance *ρ* of our statistical models for single point mutations as a function of the sum of the estimated squared bias *J_0_D* and of the variance *σ*^2^ in Eq. (3). *J*_0_(*K*) (denoted as 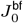 in the plot axis label) is fitted to for each value of *K* by maximizing the scaling correlation as explained in the main text. **I**: Bias factor *J*_0_(*K*) (normalized by *J*_0_(0)) inferred from maximizing the scaling correlation as in panel H.

#### Bias and variance are sufficient to explain model performance

Equation (1) stipulates that the mean squared error over fitness prediction depends on the sum of squared bias and variance of the fitness predictors. If the performance *ρ* is, in turn, controlled by this mean squared error, we expect a relation such as

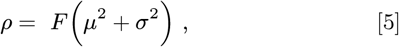

where *F* is a decreasing function of its argument.

To test the validity of Eq. (5), we compare the values of *ρ* obtained with the independent-site Potts models (*K* = 0) and different K-link Potts models (*K* = 4, 8, 16, 24) trained from various sub-alignments with different *B, D* to the sums of the squared bias and variance, see Fig. 3B. We obtain an excellent anti-correlation between *ρ* and 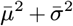 across a large range of values of *B* and *D*, in full agreement with Eq. (5) (*R* ~ 1 for every *K*–link model). The sum of squared bias and variance is by far the biggest factor in determining the predicting performance of the models.

#### Bias and variance are related to the quality and the quantity of data as predicted by theory

We then test the relation between the squared bias and the Hamming distance in Eq. (4), by generating MSAs at a given *D* and numerically computing 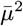 for several K-link Potts model of increasing complexity. As shown in Fig. 3C, the linear relation between the true squared bias and *D* is confirmed for every value of *K* (*R* ≃ 1 for every tested K-link model).

Similarly, we find a good agreement between the numerical variance and our theoretical estimate in Eq. (3), see Fig. 3D (*R* ≃ 1 for every K-link Potts model).

#### J_0_ *reflects the un-modeled epistasis*

The slope of the numerical bias *μ*^2^ with *D* (Fig. 3B) gives access to an estimate for *J*_0_. We plot in Fig. 3F the corresponding *J*_0_ as a function of the number *K* of links in the Potts model, from *K* = 0 (independent model) to *K* = 40. We find that *J*_0_(*K*) decreases almost linearly with *K* before reaching a saturation point around *K* = 20.

This decrease is in accordance with the notion of *J*_0_ as reflecting the un-modeled epistasis. In the context of Lattice Proteins, this saturation behavior is expected to reflect the presence of two distinct classes of un-modelled epistastic couplings. Strong pairwise interactions correspond to the *N_c_* = 28 contacts on the 3D fold (Fig. 3A). These “structural” couplings are expected to be largely responsible for the magnitude of epistatic effects in the fitness function, therefore contributing the most to the value of *J*_0_. The remaining *K_max_* – *N_c_* are weaker, and may be due to the need to avoid other folds (negative design) or to higher-order interactions (50).

To verify this hypothesis, we retrieve a pairwise approximation of the real fitness function by inferring a fully-connected Potts model from a very large alignment (*B* ~ 10^6^ sequences). We then separate the inferred Potts couplings into structural and non-structural, and compute their variance as a proxy for their expected contribution to the value of *J*_0_ (see Appendix A). As shown in Fig. 3F, structural couplings have a much larger variance than the other ones. We can devise an effective theoretical approximation of the behavior for *J*_0_(*K*) by assuming that all structural and non-structural couplings are uniformly drawn from two distributions with the two variances above, and that the sparse model progressively includes structural couplings in its energy function up to *K* = *N_c_*. The expected behavior of *J*_0_(*K*) under this effective model, shown in Fig. 3G, agrees with Fig. 3E, and saturates to its lowest value around *K* = 28, which corresponds to the total number of structural couplings.

#### J_0_ *can be inferred without any knowledge of the true fitness land-scape*

Last of all, we propose an alternative approach to estimate the bias factor *J*_0_, which is applicable to real protein data, where the sequence-to-fitness mapping is unknown. For fixed model complexity (value of *K*) we subsample the MSA, infer the corresponding K-link Potts models, and estimate the predictive performances *ρ*. The procedure is repeated by varying quantity (*B*) and quality (*D*) of the sub-MSAs. We then consider *J*_0_ as a free parameter and infer its value by maximizing the Spearman correlation between the two sides of Eq. (5), where *σ*^2^ is estimated from Eq. (3) and *μ*^2^ = *J*_0_*D*. We call this approach “best scaling fit”.

We apply this procedure to the same lattice protein data shown above. Results for the performance *ρ* vs. *J*_0_*D* + *σ*^2^ are shown in Fig. 3H for all K-link Potts models (*R* ≃ 1 for every tested K-link model), in excellent agreement with the ground truth results of Fig. 3B. The fitted values of *J*_0_(*K*) are reported in Fig. 3I, in excellent agreement with Fig. 3E,G.

### D. Performance vs. quantity and quality of sequence data for real proteins

#### Trade-off explains the predictive performance in mutagenesis experiments

The relation in Eq. (1), which we verified on in-silico proteins, postulates that the performance *ρ* of the predictive model is controlled by the sum of the squared bias *J*_0_*D*, as an inverse proxy for the the *quality* of the sequence data, and of the variance *σ*^2^, which inversely depends on the *quantity* of data. To test our theory on real data, we consider 7 different mutagenesis experiments on 7 proteins. For each protein, we sub-sample the corresponding MSA as done in Fig. 2, to obtain sub-MSAs with a large range of values of *D* and *B*, from which we can compute the estimated variance *σ*^2^. We then compute the two descriptors *D* and *σ*^2^ from each sub-MSA, and compare them with the predictive performance inferred from the data.

As reported in Fig. 4A, *D*, is a fairly good predictor for the performance of an independent-site Potts model (RNA-binding domain - absolute value of Spearman correlation coefficient *r_S_* between *D* and *ρ* = 0.70), while the variance alone correlates more weakly with the predictive performance (RNA-binding domain - absolute value of Spearman correlation coefficient *r_S_* between *σ*^2^ and *p* = 0.25). However, when the performance is compared to the sum of the squared bias and the variance, *J*_0_*D* + *σ*^2^, the correlation can be made much higher through fitting of *J*_0_ (RNA-binding domain - absolute value of Spearman correlation coefficient *r_S_* between *J*_0_*D* + *σ*^2^ and *ρ* = 0.95, Fig. 4B). This strong correlation is confirmed for the 7 protein families (all *r_S_* > 0.9 for the 7 families, Fig. 4C and Supplementary Fig. 4), providing a strong verification of the theoretical and numerical framework developed above.

**Fig. 4.**
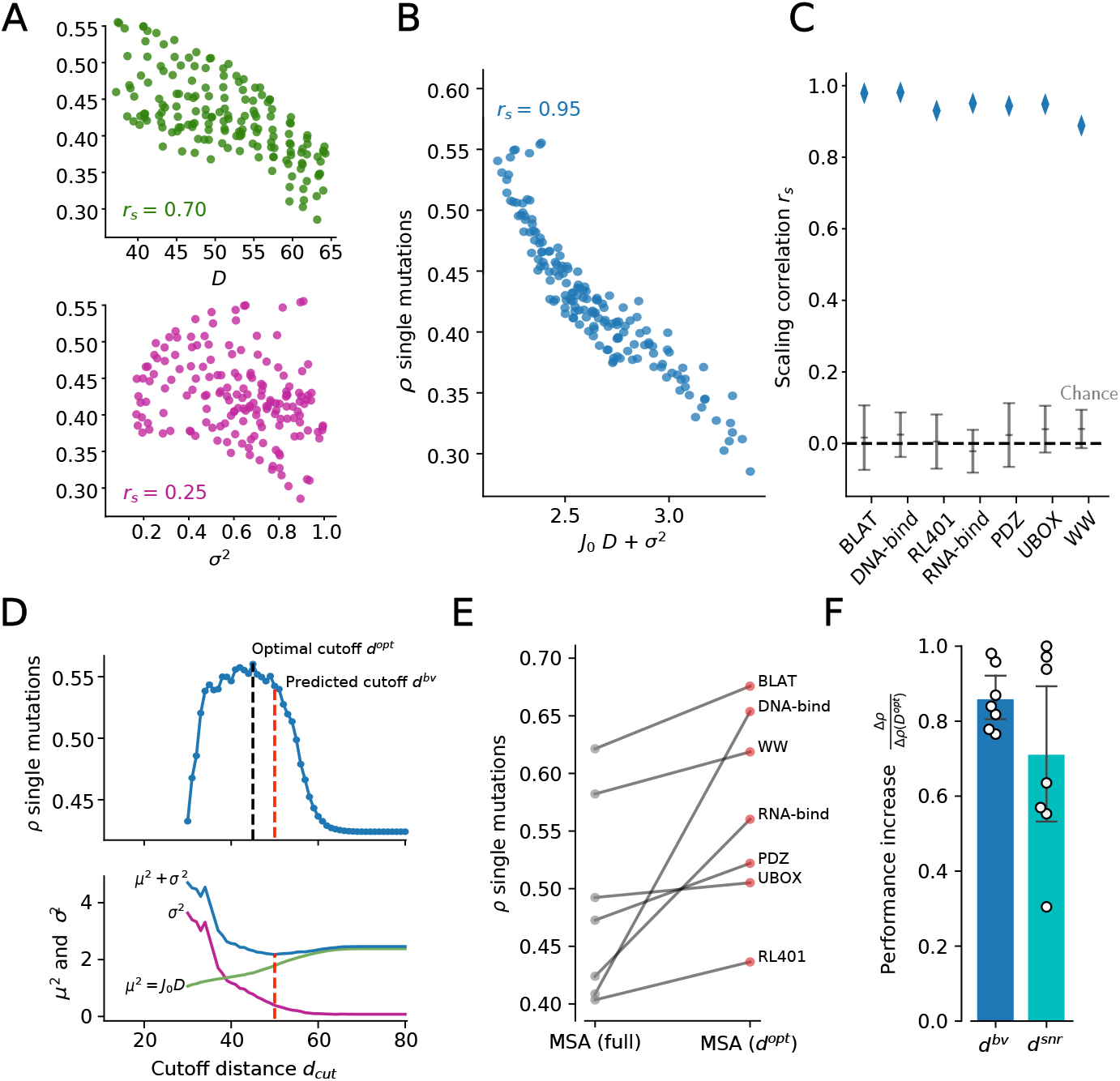
Quality-quantity trade-off explains the predictive performance of statistical modelling. **A** predictive performance of single-point mutations using the Independent-site on the RNA-bind protein, shown as a function of the mean Hamming distance of the MSA (top) and variance estimated from the alignments (bottom). **B** predictive performance of single-point mutations as a function of the linear sum of squared bias and variance. The scaling correlation *r_S_* is computed as the absolute value of the Spearman correlation coefficient of *J*_0_*D* + *σ*^2^ vs. *ρ*. The bias factor *J*_0_ is inferred by maximizing *r_S_*, as done in Fig .2 E. **C** scaling correlation *r_S_* for the seven protein families, compared to chance levels. The chance distribution is built by destroying the relationship between the performance *ρ* and the two descriptors by random order shuffling, then repeating the *J*_0_ inference procedure to account for the scaling optimization during its estimation. Error bars show standard deviations over *n* = 100 repetitions of the random shuffling. **D** top: RNA-bind family, predictive performance *ρ* as a function of the cutoff distance *d_cut_*, showing the existence of an optimal cutoff *d^opt^* (black dashed line). Bottom: individual contributions of squared bias (*J*_0_ *D*, purple line), variance (*σ*^2^, green line) and their sum (blue line). The red dashed line indicates the minimum of *J*_0_*D* + *σ*^2^, which corresponds to the predicted maximum performance cutoff *d^bv^*. **E** Values of predictive performance *ρ* at the optimal cutoffs compared to the full alignments for the 7 protein families. **F** ratio between performance increase at cutoffs of interest and at the optimal cutoff for the 7 protein families.

#### Optimization of performance through a focusing procedure

We may now exploit our understanding of how performance depend on the number *B* and on the mean Hamming distance *D* of the sequences in the MSA to find the optimal sub-alignments maximizing *ρ*.

As we see in Fig. 2, we can start from the full MSA and progressively *focus* around *wt* by excluding all sequences of “low quality”, i.e., at Hamming distances higher than a given cutoff *d_cut_*. As we lower *d_cut_* from its maximal value (*N*, number of sites) down to 0, this focusing procedure increases the variance while decreasing the bias, as we select fewer sequences with higher homology to *wt*. As already seen in Fig. 2C, the predictive performance *ρ* has a maximum at a certain optimal cutoff *d^opt^* (Fig. 4D (top panel)), highlighting the trade-off between bias and variance in controlling the performance.

In Fig. 4E we report the performance of the independent-site at the optimal cutoff *d^opt^*. We find notable improvements in the predictive performance for 6 out of 7 protein families with respect to the full MSA (mean improvement Δ*ρ*(*d^opt^*) = 0.081). Importantly, for 3 families out of 7 (DNA-bind, RL401, WW), the value or *ρ* at the optimal cutoff exceeded the best performance reported in (37) and obtained with PLM-DCA, a standard approach to learn the Potts model parameters (57). This result is striking, as the number of parameters and the number of training sequences involved in the inference at *d^opt^* are both very much reduced compared to fully-connected Potts models on large MSAs. The most outstanding illustration is DNA-bind family, where top performance (Δ*ρ* = 0.26) is found for *d^opt^* = 29, corresponding to only *B* = 37 effective sequences in the MSA (see Supplementary Fig. 5).

#### Cutoff for optimal focusing can be reliably predicted from heuristics

According to Eq. (5) the best performance is reached for the alignment that minimizes the sum *J*_0_ *D* + *σ*^2^. We call this optimal predicted cutoff *d^bv^*, as for bias-variance,

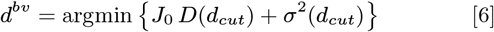

As reported in Fig. 4D(top) (red line) and Fig. 4F, this procedure allows us to predict the optimal cutoff with good precision (mean relative error = 0.08). Importantly, the performance increase at the predicted cutoff *d^bv^* captures most of the total possible improvement (mean guessed relative increase for the 7 families Δ*ρ*(*d^opt^*)/Δ*ρ*(*d^opt^*) = 0.86 ± 0.08, see Fig. 4F). Globally, the performance at the predicted cutoff *d^opt^* is systematically higher than the performance with the full MSA (mean Δ*ρ*(*d^opt^*) = 0.073, paired Wilcoxon test over the *n* = 7 families: *P* = 0.018).

However, knowledge of the bias factor *J*_0_ entering Eq. (6) is not always available, as it requires a systematic analysis of predictive performance relying on the outcome of mutagenesis experiments as a reference. We propose below a simple heuristics for predicting the optimal cutoff, requiring no experimental input and based on a signal-to-noise ratio (SNR) comparing the spread of inferred fitness values across sites and mutations and the statistical variance *σ*^2^, Fig. 4D(bottom):

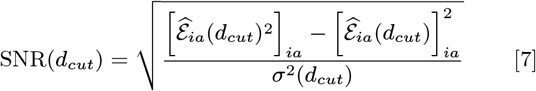

Choosing the cutoff *d^snr^* corresponding to the threshold SNR= 3, we again find systematic improvements in the predictive performance (mean guessed relative increase for the 7 families Δ*ρ*(*d^snr^*)/Δ*ρ*(*d^opt^*) = 0.71 ± 0.10, see Fig. 4F), providing an unsupervised, parameter-free criterion to select the optimal MSA for the predictive analysis.

#### The bias factor J_0_ is informative about epistatic effects not captured by the model

We repeat in Fig. 5A the approach of Fig. 4B, using the *K*–link Potts model rather than the independent-site model for fitness predictions. The number of couplings, *K*, is chosen to be a fraction of *N*, and is much smaller than *K_max_*, implying that the Potts model is very sparse. For each sub-alignment of the RNA-binding domain data we determine the best scaling fit bias *J*_0_(*K*) and We observe very high correlations between *ρ* and *J*_0_(*K*) *D* + *σ*^2^. We also observe that top performances are found for a non-zero value of *K*, e.g. *K* = 0.1 *N* in Fig. 5A. The optimal value of *K* generally varies from family to family, as reported below.

**Fig. 5.**
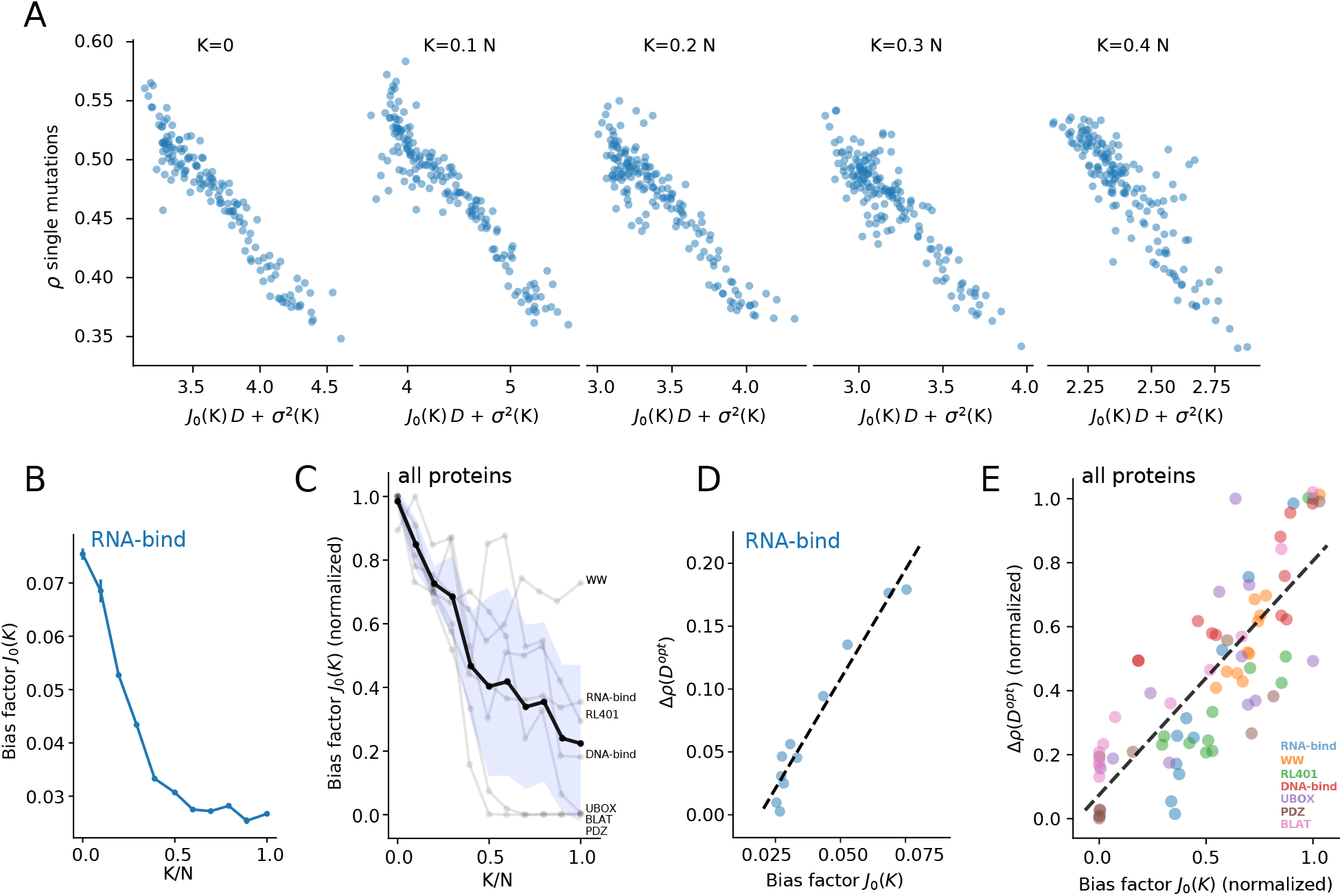
The bias factor *J*_0_ depends on the strength of un-modelled epistasis. **A** Scaling correlation between predictive performance *ρ* and *J*_0_ *D* + *σ*^2^ for the RNA-bind protein, modeled with the Sparse Potts model with different number of couplings. N is the length of the protein (82 sites). **B**: values of the bias factor *J*_0_ as a function of the number of modelled couplings in the Sparse Potts Model for the RNA-bind protein. **C**: same of B for the seven protein families combined; the black line and the blue area represent the mean and the standard deviation over the seven protein families. **D** Relation between bias factor *J*_0_(*K*) and improvement at best cutoff Δ*ρ*(*d^opt^*) for the RNA-bind protein. **E** same of D for the seven families combined. Values of *K* range from *K* = 0 to *K* = N. Each color corresponds to a different protein family as reported in the legend.

The value of the bias factor *J*_0_(*K*) is shown as a function of the number of links per site in Fig. 5B for the RNA-binding domain and for all 7 protein families in Fig. 5C. The general behaviour is similar to the one observed for lattice proteins (Fig. 3), and shows that *J*_0_(*K*) decreases with *K* until saturation is reached. As the expressive power of the predictive model increases, the squared bias decreases and is less affected by the quality of the sequence data. The saturation indicates that, above some critical *K*, adding more pairwise couplings does not help to reduce the bias. This can be interpreted as the presence of higher-order epistasis, e.g. 3-site couplings between residues, which cannot be accounted for by the *K*–link Potts model. Empirically, we expect that the focusing procedure should provide substantial improvement if the bias strongly decreases with *D*, that is, if the bias factor *J*_0_ is large, *e.g*. in the case of the independent-site model. The intuition is that, when the bias quickly decrease with the quality of the data, there is a margin for improvement of performances by removing some low-quality data, while not increasing too much the statistical variance of the inferred model parameters. We report in Fig. 5D the gain in performance *ρ* (compared to the independent Potts model, with *K* = 0) for the RNA-binding domain as a function of the bias factor *J*_0_ when *K* is varied. Results show a strong positive correlation between the two quantities. The same correlation is found across all 7 protein families, see Fig. 5E and Supplementary Fig. 6.

## Discussion

In this work, we have investigated, through a combination of analytical and numerical approaches, how the quality and quantity of sequence data determine the capability of statistical models, with variable expressive power, to predict the fitness effects of single-point mutations. As expected from the well-known bias-variance trade-off of statistics simple models require few data to be inferred, but results in systematic prediction errors (bias *μ*). Conversely, powerful models are in principle capable of expressing complex sequence-to-fitness relationships but their many defining parameters are subject to more statistical errors due to the limited amount of available data (variance *σ*^2^). We have shown that a good predictor of performances was given by the sum *μ*^2^ + *σ*^2^, and have analytically related the variance to the number of sequences *B* in the alignment and the squared bias *μ*^2^ to the evolutionary depth, estimated through the mean Hamming distance *D* to the mutated wild type sequence. Our theory was quantitatively confirmed by extensive tests on *in silico* lattice proteins for which the ground-truth fitness is known, and on mutagenesis datasets of 7 proteins families we have analyzed. Based on the results above, we then proposed a “focusing” procedure to optimally select the best subset of sequences from a multi-sequence alignment, and tested it on the 7 mutagenesis experiments. With this procedure, the least powerful, independent sites model, showed performances higher than fully connected graphical models trained on the same data for 4 out of 7 studied protein families, and comparable performances for the remaining ones.

An important finding of the present work is the so-called bias factor *J*_0_, which relates the squared bias *μ*^2^ to the mean Hamming distance *D* of the sequence data to the wild-type sequence: *μ*^2^ ≃ *J*_0_ *D*. Importantly, the analytical derivation of this relation showed that *J*_0_ accounted for un-modelled epistasis, i.e., for the statistical properties of the fitness landscape that cannot be reproduced by the class of models considered. This result has two consequences, both conceptual and practical. First, it explicitly demonstrates that key information about the amount of epistasis in fitness landscapes is, in principle, accessible even with models with limited complexity, constrained by data availability. From a practical point of view *J*_0_ can be estimated through a regression of the performance *ρ* vs. a linear combination of *D* –chosen at will through subsampling of the multi-sequence alignment– and *σ*^2^ –given by Eq. (3)–, see Fig. 5A; this procedure can therefore be applied to any protein family, for which sequence and mutagenesis data are available. Second, the meaning of *J*_0_ emphasizes the role of the expressive power of the model in the relative importance of the bias and variance terms, and to what extent each one of these factors affect performances. The value of *J*_0_ is a good predictor of how much can be gained in performance by pruning the sequence data and focusing around the *wt* sequence (Fig. 5E). This result entails that simpler models have higher potential for improvement in fitting a local neighborhood through focusing, and can overcome complex models when training data is appropriately selected.

Determining the optimal cutoff distance for focusing can theoretically be done following the quantity-quality trade-off analysis presented above, *e.g*. using some already available mutagenesis experiment. We proposed an empirical rule that did not require any mutational information and was based on a signal-to-noise criterion. This empirical cutoff led to systematic improvement of performance for all tested families (Supplementary Fig. 5).

Our focusing and modeling procedures could be further improved along several directions. First, in the the *K*-links Potts model considered here we have selected relevant links according to the Frobenius norms of the couplings of the inferred Potts model (equivalently, in the Direct Coupling Analysis, DCA). The rationale for this criterion is that the coupling norm is a good proxy for coevolution and contact between residues. Sparsity of the interaction graph can be enforced, within DCA, through *L*_1_ regularization over the couplings (54, 58, 59). However, in a related work (38), we have shown that DCA-based ranking is not an optimal predictor of relevance of couplings for protein function. Couplings can be better selected using a semi-supervised procedure, which exploits a subset of mutational data. Such optimally selected *K*-Links Potts models achieve a clear increase of the performances in predicting the effect of mutations.

Second, we have estimated, so far, the closeness of an alignment to the wild-type sequence through the average Hamming distance *D*. This choice is justified both by its simplicity, and the deep relation between *D* and the (squared) bias. However it would be worth considering more refined estimates for the distances, taking into account the phylogeny of the sequence data. Residue conservation can be assessed according to mutational history (30, 31), or to their relevance in the functionality under consideration. In addition, our focusing procedure could make use of alignment methods based on local homology, recently used to discover specific functionality proper to some protein subfamilies (60).

Our theoretical study could help improve models and alignment processing for predicting the effects of missense mutations and their impact in genetic diseases (6–10). Natural alignments of sampled missense mutations are limited in depth and naturally focused around the human genome, making independent-site models (or *K*-link Potts models with small *K* values) especially adequate. It would be very interesting to apply our focusing approach to understand how to best select sequence data in this context.

The capability of deriving optimal independent-site models, whose parameters are tuned according to the region in the sequence space under focus, could be also be important for phylogeny studies. Inferring phylogenetic relations between a set of sequences requires the capability to compute transition probabilities under a mutation-selection process. Independent-site models are particularly attractive in this regard as they lead to mathematically tractable expressions for the transitions (61), but cannot describe complex sequence-to-fitness mappings. An alternative would be to use multiple focused independent-site models to compute transitions, adapted to the multiple portions of the sequence space explored by the phylogenetic tree.

Last of all, we stress that the question of how to select the best subset of data ensuring optimal performance given a statistical model is of general interest for machine learning. Our focusing procedure is conceptually related to local regression methods, such as moving least squares approaches, which aim at locally fitting a function from data. It is therefore expected that it will find applications beyond the prediction of fitness considered here. For instance, focused independent-site models could be useful in the context of gene expression, where microarray data generally suffer from missing values, impeding the use of many multivariate statistical methods (62).

## Materials and Methods

### Multiple sequence alignments

Proteins families and the corresponding alignments were taken from (37). The alignment procedure of EVmutation (https://github.com/debbiemarkslab/EVmutation) is based on a query against the UniRef100 database of nonredundant protein sequences (release 11/2015) (63) from the wild-type sequence, using the profile HMM homology search tool jackhmmer (64) and choosing a default score in the alignment depth of about 0.5 bits/residue; the threshold was adjusted if the alignment had not enough coverage or number of sequences (37). Redundant sequences were removed from the alignments, as well as sites with more than 50% of gaps in the alignment. The list of families and wild type, the sequence length, the number of sequences, and the reference to the mutational scans are given in Table 1. Alignments are made available in Supplementary Information.

**Table 1.** From left to right: Numbers *N* of sites, *B* of sequences (after removal of redundant sequences from the alignment), 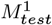 of tested single mutations, *M*^1^ of possible single mutations, and corresponding references.

| Protein Name (*wt* name) | $N$ | $B$ | $M_{test}^1$ | $M^1$ | Ref |
| --- | --- | --- | --- | --- | --- |
| Betalactamase (BLAT-ECOLX) | 253 | 7620 | 4610 | 4807 | (19) |
| DNA Binding domain (GAL4-YEAST) | 63 | 11278 | 1196 | 1197 | (65) |
| Ubiquitin (RL401_YEAST) | 71 | 10023 fit | 1160 | 1349 | (22) |
| RNA Binding (PABP-YEAST) | 82 | 70779 | 1188 | 1558 | (14) |
| PDZ (DLG4-RAT) | 84 | 24795 | 1577 | 1596 | (13) |
| UBOX Domain (UBE48-MOUSE) | 76 | 6153 | 614 | 1444 | (20) |
| WW (YAP1-HUMAN) | 31 | 8251 | 319 | 589 | (21) |

### Sequence re-weighting and MSA descriptors

We partially corrected for sampling biases by using a re-weighting procedure with 80% homology threshold in all statistical estimates on sequence data (32). We therefore define the weight of a sequence *s* to be

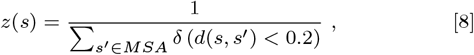

where *δ*(*X*) = 1 if condition *X* is satisfied, and 0 otherwise,and *d*(*s, s*’) is the normalized Hamming distance between sequences *s* and *s*’. The MSA descriptors were then computed as weighted averages:

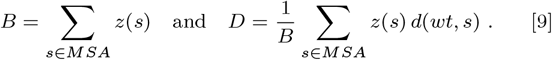

### Fitness predictions and comparison with experiments

With the *K*-sparse Potts model the probability of the sequence *s* reads

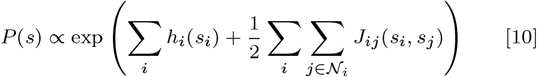

up to a normalization constant. 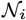 denotes the set of sites connected to site *i* on the interaction graph. The predicted fitness difference is defined as the difference in the log probabilities of the wild-type sequence (*wt*) and of the one where the amino acid at site *i* in is substituted with the amino acid *a* as *wt_i→a_*:

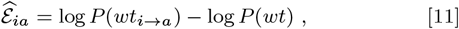

giving Eq. (2). The predictive performance *ρ* of the model is then computed as the Spearman rank correlation between experimental measures of delta-fitness for single-point mutations and the corresponding predictions.

### Inference of sparse Potts models

Following a number of recent works (34, 36–38, 58), we predicted the effects of single point mutations by inferring a Potts model from sequence data in the alignment. Here, we employed a K-link Potts model introduced in (38), where we constrain the model to have non-zero couplings only on *K* statistically-relevant links (*i, j*) (*K* = 0 being the independent model, *K* = *N*(*N* – 1)/2 the fully connected Potts model).

We chose the *K* links by scoring each link (*i, j*) as done for contact prediction in DCA analysis: we inferred a fully-connected Potts model with parameters optimized to perform contact prediction by pseudo-likelihood maximization (57); from the resulting couplings ***J**^PLM^* we defined a score for each link (*i, j*) based on the Frobenius norm of the two-sites coupling matrix 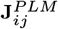. Finally, we selected those *K* pairs (*i, j*) that showed the highest Frobenius score. We then used a two-site approximation to re-infer the value of the ***J**_ij_* matrix for each of these *K* pairs given the sparsity constraint (38).

### Sub-sampling the MSA allows for varying data quality and quantity

To create new MSAs of different degrees of quality and quantity for real protein families, we sub-sampled the corresponding MSA using the following procedure. We first chose a target Hamming distance *D* and a number of sequences *B*_0_ (before re-weighting). We then randomly sampled *B*_0_ sequences *s* from the full MSA (without repetition) with a probability decreasing with the Hamming distances *d*(*s, wt*) between the sequences *s* and the wildtype:

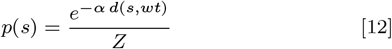

where *Z* = ∑_s′_ ∈ *MSA* ^*e*−α*d*(*s*′, *wt*)^. The parameter *α* was optimized to reach the defined *D* within a given precision (here set to 0.01). From each sampled sub-MSA, we then computed the effective number of sequences *B* as described above, as well as the variance *σ*^2^ for each sparse Potts model with *K* links through Eq. (3).

We repeated this procedure for all combinations of 16 values of *D* ∈ [0.4*N*, 0.8*N*], where *N* is the protein length, and 10 values of *B*_0_ in a range that depended on the protein family and its initial MSA size. Doing so, we obtained a population of 160 sub-MSAs with as many corresponding values of *D* and *B*.

### Lattice proteins

Lattice proteins are *in silico* proteins of fixed sequence length (*N* = 27) folding on the sites of a 3 × 3 × 3 cube (47, 49, 55). The protein *family* attached to a specific fold *F* is defined as the set of sequences *s* with low (favourable) folding energies *ϵ*(*F, s*) in *F* and unfavourable folding energies *ϵ*(*F*’, *s*) for all other possible folding structures *F*’ (little competition) (56); *ϵ*(*F, s*) is defined as the sum of Miyazawa-Jernigan interactions (66, 67) between residues *s_i_, s_j_* in contact on structure *F*. The fitness of a protein *s* (with respect to the native fold *F*) is defined as

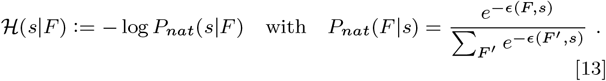

An MSA for the family *F* can then be obtained by sampling from the effective Hamiltonian 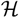. To control for the mean Hamming distance from a given wildtype sequence *wt* of the sampled MSA, we follow the procedure of (50) and sample from a biased Hamiltonian

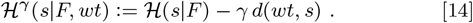

As in (50), Monte Carlo sampling is performed with Metropolis rule at effective temperature *β* = 1000 and with *T* = 1000 thermalization steps between each sampled sequence. Precise values of *D* were obtained by sub-sampling and mixing four large alignments obtained with *γ* = 0, 0.025, 0.050, and 0.075. From each MSA, the computation of descriptors *σ*^2^ and *D* as well as training and performance assessment of Potts models were performed as explained below for real proteins, with the difference that no re-weighting procedure was applied to lattice proteins data.

### Numerical estimation of bias and variance in Lattice Proteins

In the case of Lattice Proteins, we numerically computed the real fitness difference caused by single-point mutations as 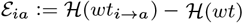. For a given inferred Potts model, we then computed the bias 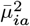 and variance 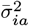 of its delta-fitness predictors 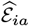 as

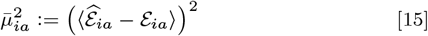

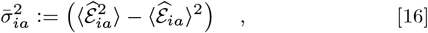

where averages are computed over *n* =10 inferences performed on as many sampled alignments with fixed number *B* of sequences and mean Hamming distance *D* to *wt*.

To relate these quantities to the single predictive performance value *ρ* of the inferred model, we defined two global measures that account for all single-point mutations (*i, a*):

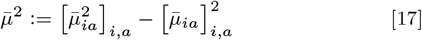

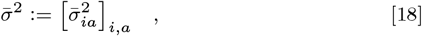

where [·]_*i, a*_ denotes the averages over sites and mutations, and the global shift in the bias 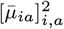 removed, as the Spearman rank correlation *ρ* is invariant under the addition of a constant to 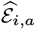. In these numerical settings, some mutations are so deleterious that will never be observed in the data, and their effect is systematically estimated by regularization only. To avoid that these outliers dominate the averages above, we restricted our analysis to those mutations that satisfy an “observability” criterion of 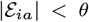. Unless specified differently, we use *θ* = 5.0 throughout all Lattice Protein results.

## ACKNOWLEDGMENTS

The authors are grateful to J. Tubiana and M. Molari for insightful discussions, and to J. Fernandez de Cossio Diaz and E. Mauri for a careful reading of the manuscript.

## Supplementary Information

### Appendix A: approximate expression of the variance

Following (38, 54) we write an approximate expression for the variance 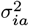 based on the so-called 2-site approximation for the biases and couplings appearing in the *K*-links Potts model:

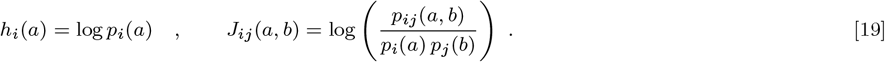

These parameter values are exact for sparse interaction graphs with a tree-like structure.

Inserting Eq. (19) into Eq. (2) we obtain

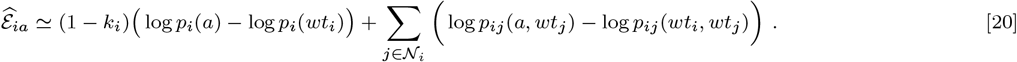

Each frequency *p* is stochastic as it varies with the sequence data, *i.e*. the sub-MSA. Approximating the variance var(*p*) = 〈*p*〉(1 – 〈*p*〉)/*B* ≃ 〈*p*〉/*B*, where 〈〉 denotes the mean over repetitions of the sub-sampling procedure, we obtain the following expression for the variance of 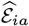,

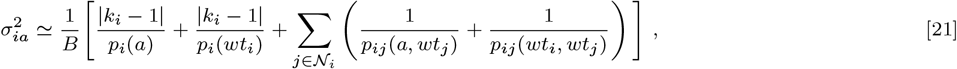

to the leading order in 1/*B*. The presence of the absolute values | · | ensures the validity of the formula for non-interacting sites *i*, such that *k_i_* = 0. Averaging Eq. (21) over the sites *i* and the mutations *a* yields Eq. (3) of the main text. Throughout this work, we considered only those mutations (*i, a*) that have at least one occurrence in the MSA to compute this average.

### Appendix B: dependence of the squared bias on the mean Hamming distance to the wt sequence

Estimating the bias of a statistical model is generally complicated since it requires knowledge of the ground-truth probability distribution that generated the data. However, we show below that, under some simplifying hypothesis, the bias can be related to the mean Hamming distance between the sequences in the MSA and the wild-type sequence, *wt*.

This appendix is organized as follows. We first define the grounth-truth distribution of sequences, that is, the fitness landscape for sequences and estimate the effect of a mutation to *wt*. We then present the predictor of this mutational effect corresponding to the independent-site model, and derive a general formula for the bias that involves the amino-acid statistics in the MSA. We then estimate how these statistical quantities depends on the MSA properties in a simple probabilistic framework for generating MSA. Last of all, we discuss how these results are changed when the model used to predict mutational effects include epistatic couplings.

#### Change in log probability following a mutation

We assume that the protein family under consideration is defined by a distribution *P* over the space of sequences *s*. Informally speaking, good sequences *s, i.e*. corresponding to functional proteins have large values of *P*, while bad sequences correspond to very low values; We further assume that the landscape associated to *P* includes local biases acting on residues as well as pairwise epistatic interactions. More precisely, we have

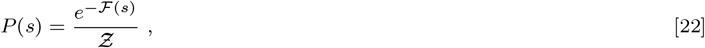

where 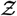 is a normalization constant, and the “statistical energy” 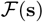 reads

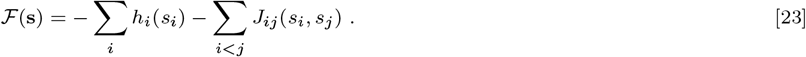

According to Eq. (23), under the mutation *wt_i_* → *a*, the variation in the log probability of the sequence is equal to

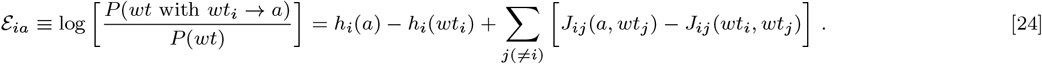

In the following, 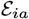 defined in the above equation will be our ground-truth value for minus the fitness of the mutated sequence relative to 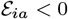 corresponds to beneficial mutations, while 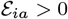 signals a deleterious mutation.

We remark that the distribution of the sequences in Eq. (22) is overparametrized. No change of the local biases of the form *h_i_*(*s*) → *h_i_*(*s*)+*b_i_* will affect *P*, neither will changes to the couplings of the form *J_ij_* (*s, s*′) → *J_ij_* (*s, s*′) + *C_ij_* (*s*) + *d_ij_* (*s*′). We may therefore, without any loss in generality, impose that all local fields and all couplings vanish when one of their amino acids coincide with the residue in the wt sequence, *i.e*. *h_i_*(*wt_i_*) = *J_ij_*(*wt_i_, s*) = *J_ij_*(*s, wt_j_*) = 0 for all residues *s* (34). With this particular choice of gauge, hereafter referred to as *wt* gauge, the expression for the change in log probability following the mutation simplifies into

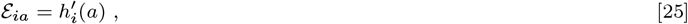

where ′ indicates the *wt* gauge.

#### Mean-field expression of bias for the independent-site model

Let us assume that we have generated a set of *B* sequences, called MSA, from the distribution defined in Eq. (22). We now want to build a model from this sequence data to have predictors of the mutational effects, 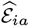. In the simplest model, residues attached to different site are independent, and the probability of a sequence is simply given by

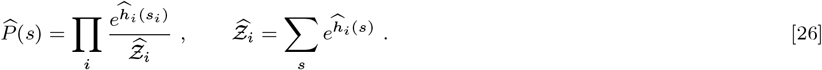

where the local fields (PWM) 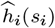 are inferred to reproduce the statistics of residues in the MSA. Choosing again the *wt* gauge, that is, setting the field values for *s_i_* = *wt_i_* to zero for all sites *i*, we have

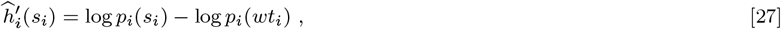

where *p_i_*(*s*) is the frequency of amino acid *s* on site *i* in the data.

According to the independent-site model,the variation in the log probability of the sequence following the mutation *wt_i_* → *a* is equal to

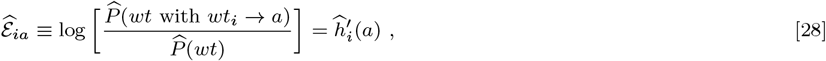

where ′ refers again to the *wt* gauge. The bias is therefore given by

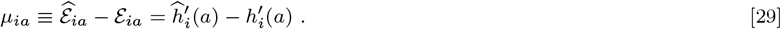

We now need to estimate this bias, more precisely, to compute 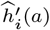 from the ground-truth distribution in Eq. (22) defined by the parameters *h_i_, J_ij_*. To make this calculation tractable we resort to the so-called mean field approximation of statistical mechanics. According to mean-field theory the effective field 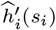 acting on a site can be approximated as the sum of the local field, *h_i_*(*s_i_*), and of the action of the other sites it is coupled to, substituted with their mean occupancies. More precisely,

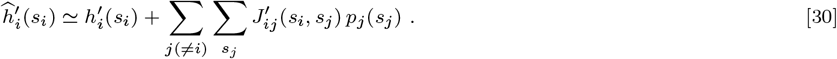

As a conclusion, based on Eq. (29), our mean-field expression for the bias in predicting the effect of mutation *a* on site *i* is equal to

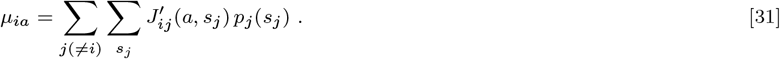

Two remarks are important here. First the bias vanishes when the couplings are equal to zero. Indeed, if the ground-truth distribution of amino acids factorizes over sites, then there is no systematic loss of accuracy in inferring the distribution with an independent-site model; statistical errors in inferring the fields from the MSA data will contribute to the variance term in the bias-variance trade-off, but the bias vanishes. Second, due to the choice of the *wt* gauge the coupling *J*(*a, s_j_*) vanish when *s_j_* = *wt_j_*. We may therefore rewrite the bias as

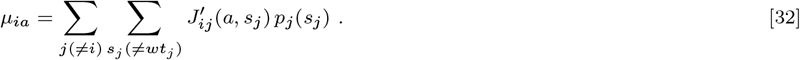

#### Relationship between the squared bias and the statistics of residues in the MSA

As our measure of goodness of prediction relies on the Spearman correlation between the experimental and predicted changes in fitness, respectively, Δ*E_ia_* and 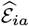, it is left unchanged by any additive constant to 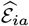. We should therefore consider the centered bias

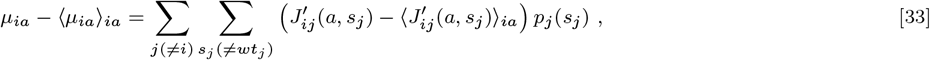

where 〈·〉_*ia*_ denotes the average over all sites *i* and residues *a* (different from the *wt* residue).

While we do not know the ground-truth fitness landscape from which sequences are drawn we may use DCA estimates of local fields and couplings as proxies for the true *h_i_* and *J_ij_*. In all cases studied in this work we find that the average coupling is very small compared to the standard deviation *J*_0_, and can be replaced with zero. The centered bias is therefore practically equal to the bias.

We may now estimate the average squared bias over all sites and mutations

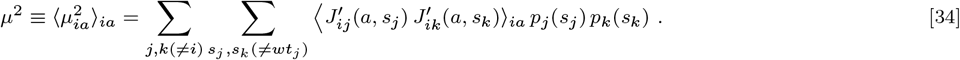

Neglecting correlations between the couplings on different sites only pairs of identical sites *j* = *k* carrying identical amino acids *s_i_* = *s_k_* contribute to the sum above, with the result

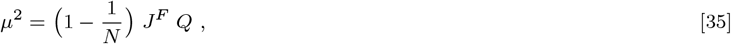

where *J^F^* is the variance of the couplings, *N* the length of the protein, and

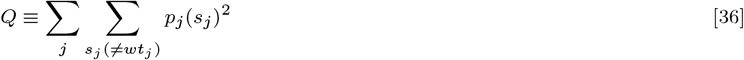

is a characterization of the MSA statistics. The validity of this formula is confirmed by studies of lattice-protein models, for which the bias can be calculated exactly, see Supplementary Fig. 1.

We show in the next subsections that the average squared frequency *Q* is, to a very good approximation, proportional to the mean Hamming distance of the sequences in the MSA to the *wt*. This result justifies the linear dependence of the squared bias *μ*^2^ upon *D* reported in the main text.

#### Statistical properties of the subsampled MSA

Consider a multi-sequence alignement (MSA) with *B_tot_* sequences of length *N*. For the sake of mathematical tractability, we assume that all sequences are drawn from a background (*bg*) distribution, where the probability of the amino acid *a* is denoted by *bg*(*a*), see for instance Carugo, O. (2008), Amino acid composition and protein dimension. Protein Science, 17: 2187-2191. https://doi.org/10.1110/ps.037762.108. We now build a sub-MSA as follows:

- One of the sequences is the MSA is called *wt* and included in the sub MSA;
- Each other sequence in the MSA is retained with probability ∝ *e^α d^*, where *d* is its Hamming distance to *wt*, and *α* is a positive parameter.

**Supplementary Figure 1.**
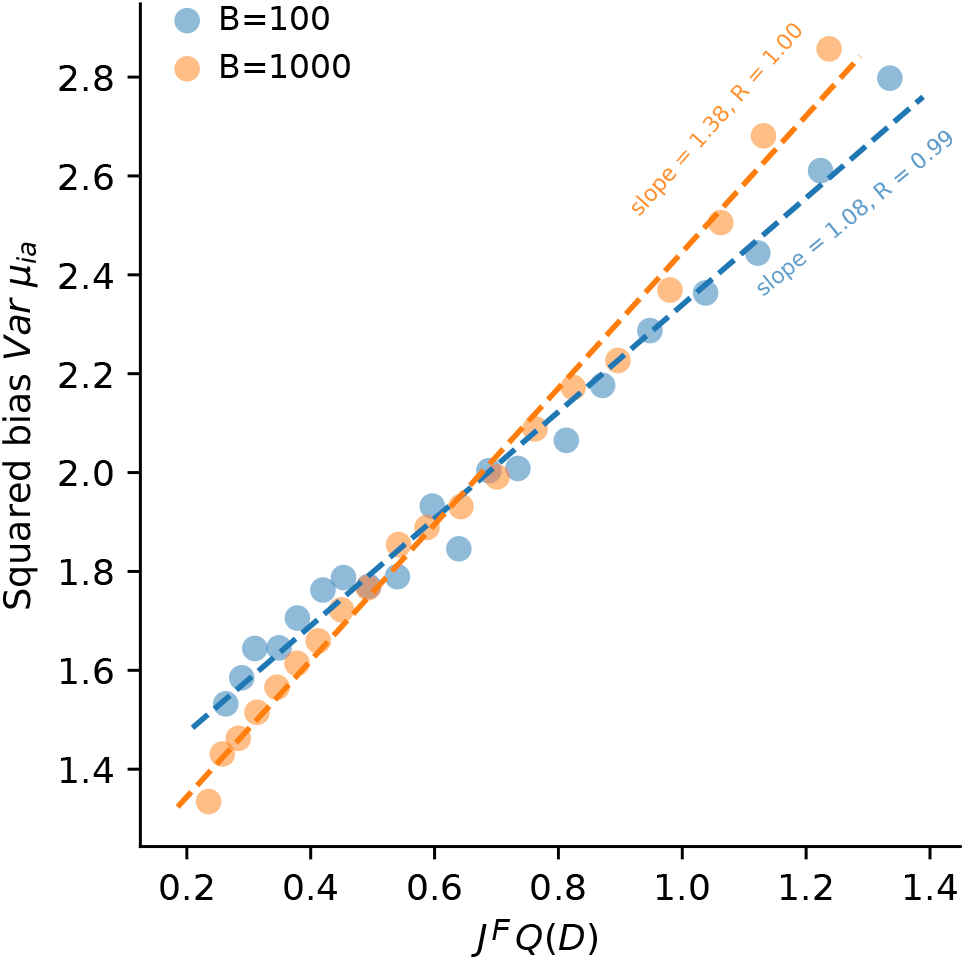
Squared centered bias from Eq. (33) vs. *J^F^* × *Q*, where *J^F^* is estimated as the variance of the couplings and *Q* is defined in Eq. (36), for the lattice-protein model associated to the structure shown in Main text Fig. 3A.

We want to estimate, as functions of *α*: (1) the mean Hamming distance of the sequences in the sub-MSA to *wt*, *D*_*sub*–*MSA*_; (2) the mean number of sequences in the sub-MSA, *B*_*sub*–*MSA*_; (3) the average squared frequencies, *Q*_*sub*–*MSA*_, entering eqn (36).

We first focus on all sequences in the sub-MSA but *wt*. According to the sub-sampling procedure above, the probability of amino acid *a* on site *i* is proportional to *bg*(*a*) if *a* = *wt_i_* and to *e*^−*α*^ *bg*(*a*) if *a* ≠ *wt_i_*. The mean Hamming distance of sequences to *wt* is therefore

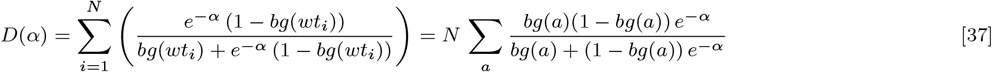

where the last equality comes from the average over the *wt* sequence. Similarly, we get

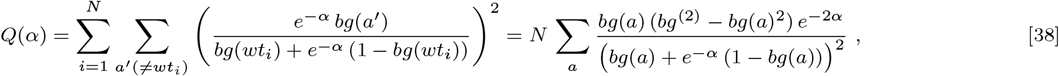

where, for integer-valued *k*,

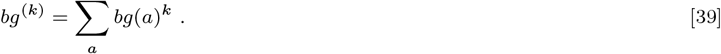

To estimate the number of sequences that are subsampled, we consider the generating function of distances,

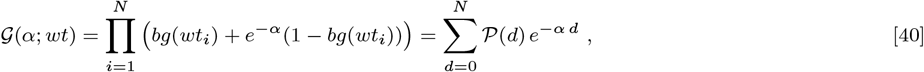

where 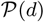 is the probability that a sequence is at distance *d* from *wt*. The average number of sub-sampled sequences can therefore be approximated as

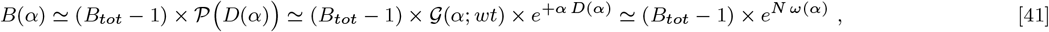

where

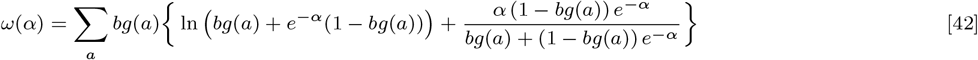

after averaging of 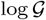 over the *wt* sequence.

Inserting back the *wt* sequence in the sub-MSA, we obtain the following expressions for the three quantities of interest:

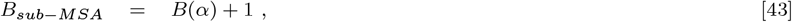

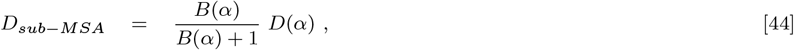

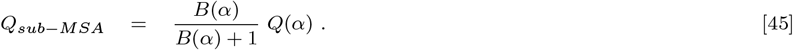

#### Approximate linear dependence of *Q* upon *D*

We now parametrically plot the number *B*_*sub*–*MSA*_ of sequences in the sub-MSA and the averaged squared frequencies *Q*_*sub*–*MSA*_ vs. the average Hamming distance *D*_*sub*–*MSA*_ by varying *α* from 0 (all *B_tot_* sequences are considered) to +∞ (all sequences are left out, with the exception of *wt*. Results are shown in Supplementary Fig. 2. We observe that *Q* is approximately a linear function of the distance *D* over a large range of sub-sampling levels of the full MSA.

**Supplementary Figure 2.**
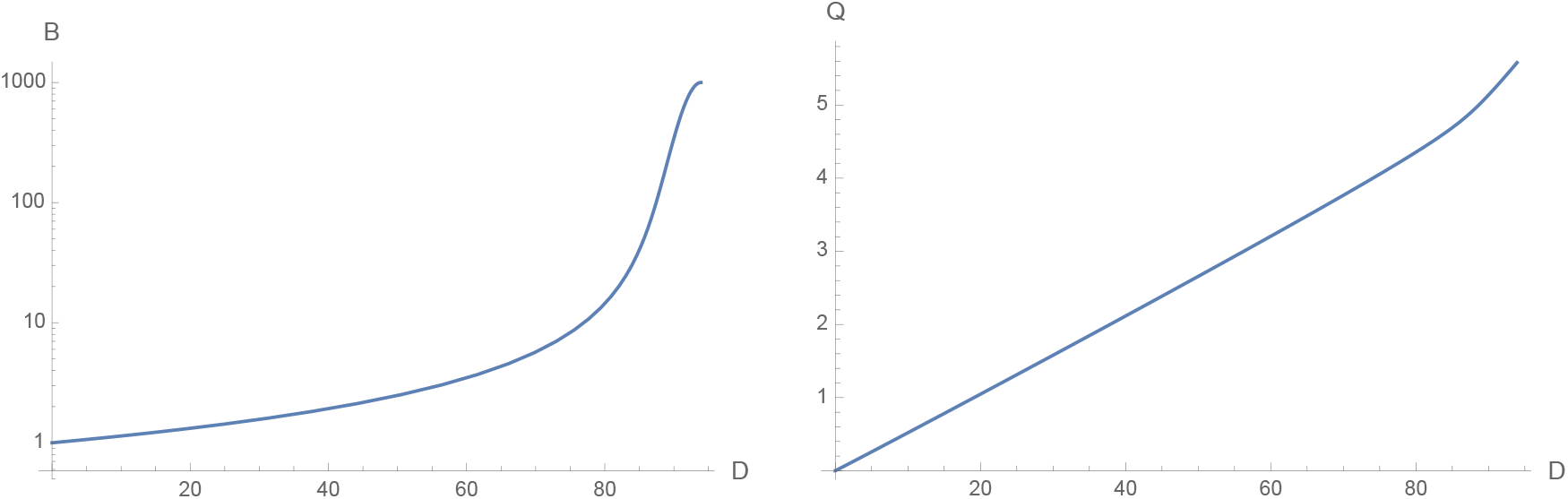
Number *B*_*sub*–*MSA*_ of sequences in the sub-MSA (left) and averaged squared frequencies *Q*_*sub*–*MSA*_ (right) vs. average Hamming distance *D*_*sub*–*MSA*_, obtained from eqns. (43), by subsampling a MSA with *B_tot_* = 1000 sequences of length *N* = 100. The approximate linear scaling of *Q* with *D* is valid even when the full MSA is very strongly subsampled. The parts of the curves corresponding to distances *D* ≤ *D_min_* ≃ 40 include on average less than 2 sequences in the sub-MSA (hence a random realization of the sub-MSA may contain the *wt* only), and should be discarded, see text.

An estimate of the slope *β* can be computed through an expansion of *ω*(*α*) in eqn (42) to the second order in *α*. We obtain

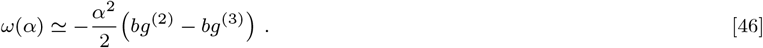

Based on this quadratic approximation, we conclude:

- For *α* = 0, the full MSA is retained through sub-sampling, with

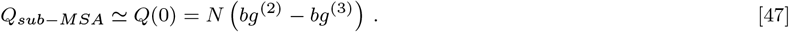 The average distance to *wt* is

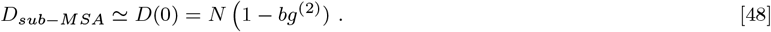
- As *α* increases, the sub-MSA shrinks until it contains *wt* only. This happens for *α* = *α_min_* such that

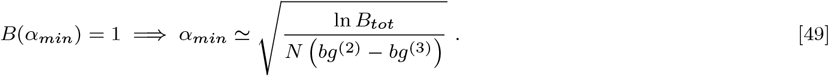 We see that *α_min_* is small, and the quadratic expansion accurate for long sequences or small MSA.
- For larger values of *α*, the predictions of eqn (43) are meaningless since they are based on sub-MSA with, on average, less than 2 sequences, see Supplementary Fig. 2. In practice, most sub-MSA contain *wt* only, and *Q*_*sub*–*MSA*_ = 0.

We can therefore estimate the slope through

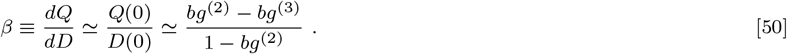

The above expressions were obtained with a uniform background model along the sequence, but can easily be extended to the case of *bg*(*a*) varying from site to site.

#### Case of non-homogeneous pairwise couplings: inference with the sparse-Potts model

Formula (35) above implicitly assumes that all couplings *J_ij_*(*a, b*) have the same variance *J*_0_, independently of the sites *i, j* under consideration. In reality, however, the couplings entering the ‘ground-truth’ statistical energy 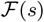 (23) may considerably vary with the pairs of sites *i, j*. This situation may in particular arise when the sites *i, j* are in contact on the protein fold.

For the sake of simplicity, let us assume that

- there are *M*_+_ pairs of sites *i, j* such that the variance of the couplings (computed over the amino acids carried by the sites) is

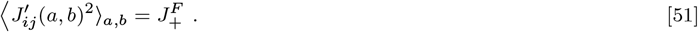
- for the remaining 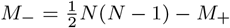 pairs of sites *i, j* the variance of the couplings is

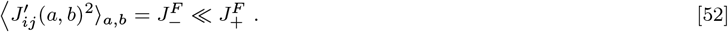

Assume we now predict the mutational effect with the sparse Potts model, with *K* couplings allowed to take non-zero values. Equation (22) for the probability of a sequence is therefore substituted with

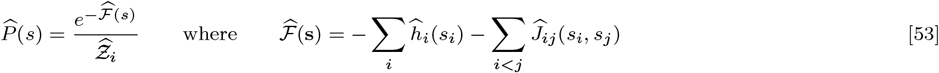

and 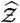 is a normalizing factor.

The calculation of the predicted change in log probability following a mutation is straightforward from the previous considerations, and eqn (29) for the bias is still valid. However, upon application of mean-field theory to both the inferred sparse Potts model and the ground-truth distribution, eqn (30) becomes

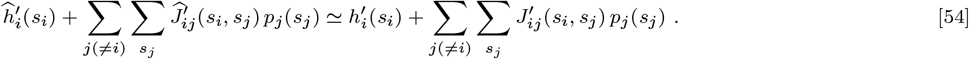

We may conclude that the bias is now equal to

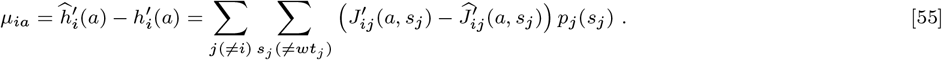

***Case*** *K* ≤ *M*_+_. If the Potts model used to reproduce the fitness landscape is very sparse, the *K* inferred couplings 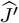 are likely to be good approximations of the *K* largest ground-truth epistatic couplings *J^′^* (having variance 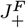). As a consequence, *K* terms in the sum in eqn (55) vanish. After averaging over the sites and amino acids, the squared bias *μ*^2^ is given by

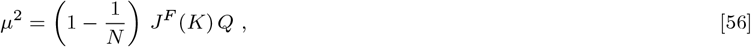

where *Q* is given by eqn (36), and

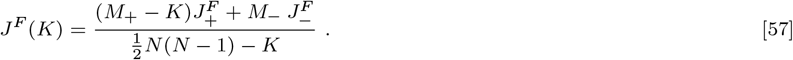

Notice that the effective coupling variance *J^F^*(*K*) is a decreasing function of *K*: as more and more couplings are included in the inferred Potts model less and less epistatic effects remain un-modelled and contribute to the bias.

***Case*** *K* > *M*_+_. Equation (56) still holds, with

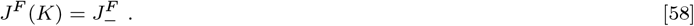

In this regime, the effective coupling variance does not vary with *K* any longer.

## Supplementary figures

**Supplementary Figure 3.**
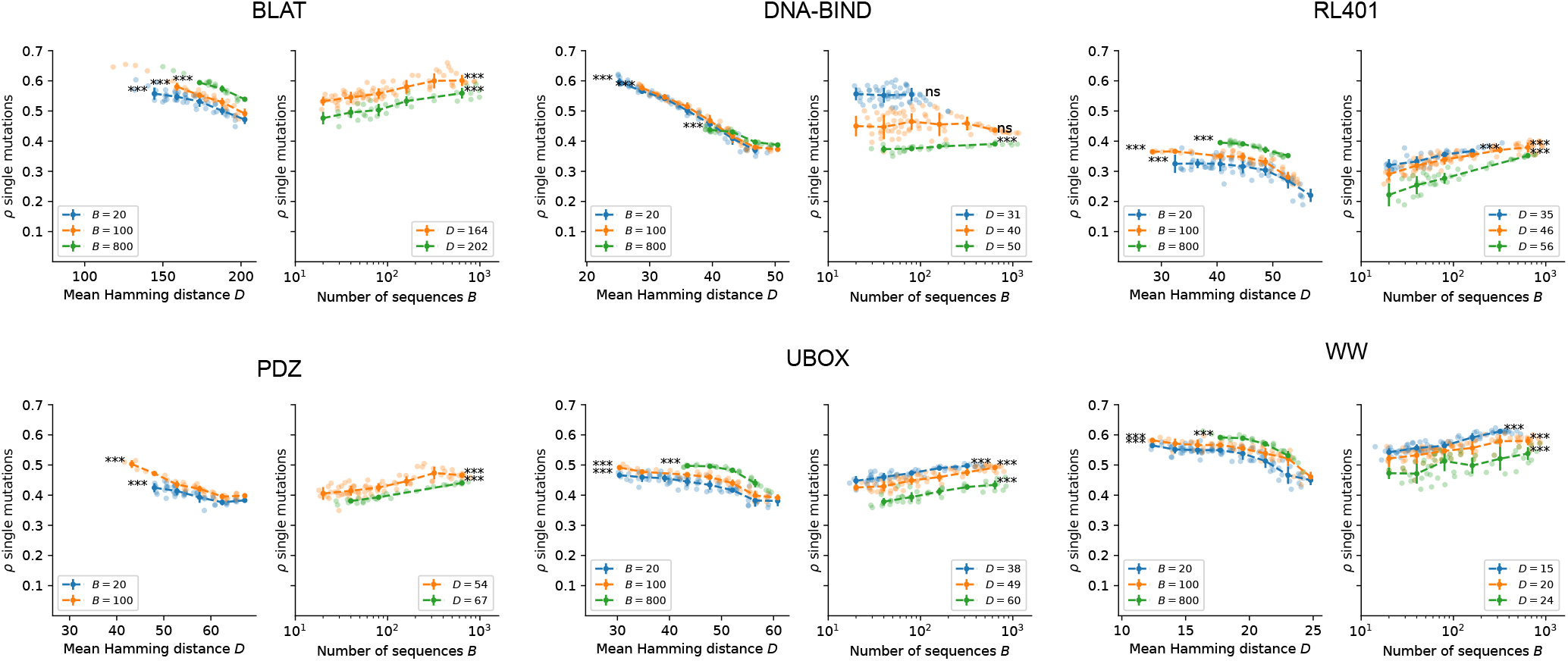
Same as Main text Fig. 2D for all protein families except RNA-Bind (shown in Main text Fig. 2D): systematic analysis of the predictive power *ρ* as a function of the mean Hamming distance *D* of sub-alignments with fixed size *B* (left panels), and of the sub-alignment size *B* at fixed Hamming distance *D* (right panels). Each point represents the binned average and standard deviation of several sub-samples obtained at the corresponding values of *D* and *B* (see Methods). All significance levels refer to Spearman rank correlation. * P<0.05; ** P<0.01; *** P<0.001.

**Supplementary Figure 4.**
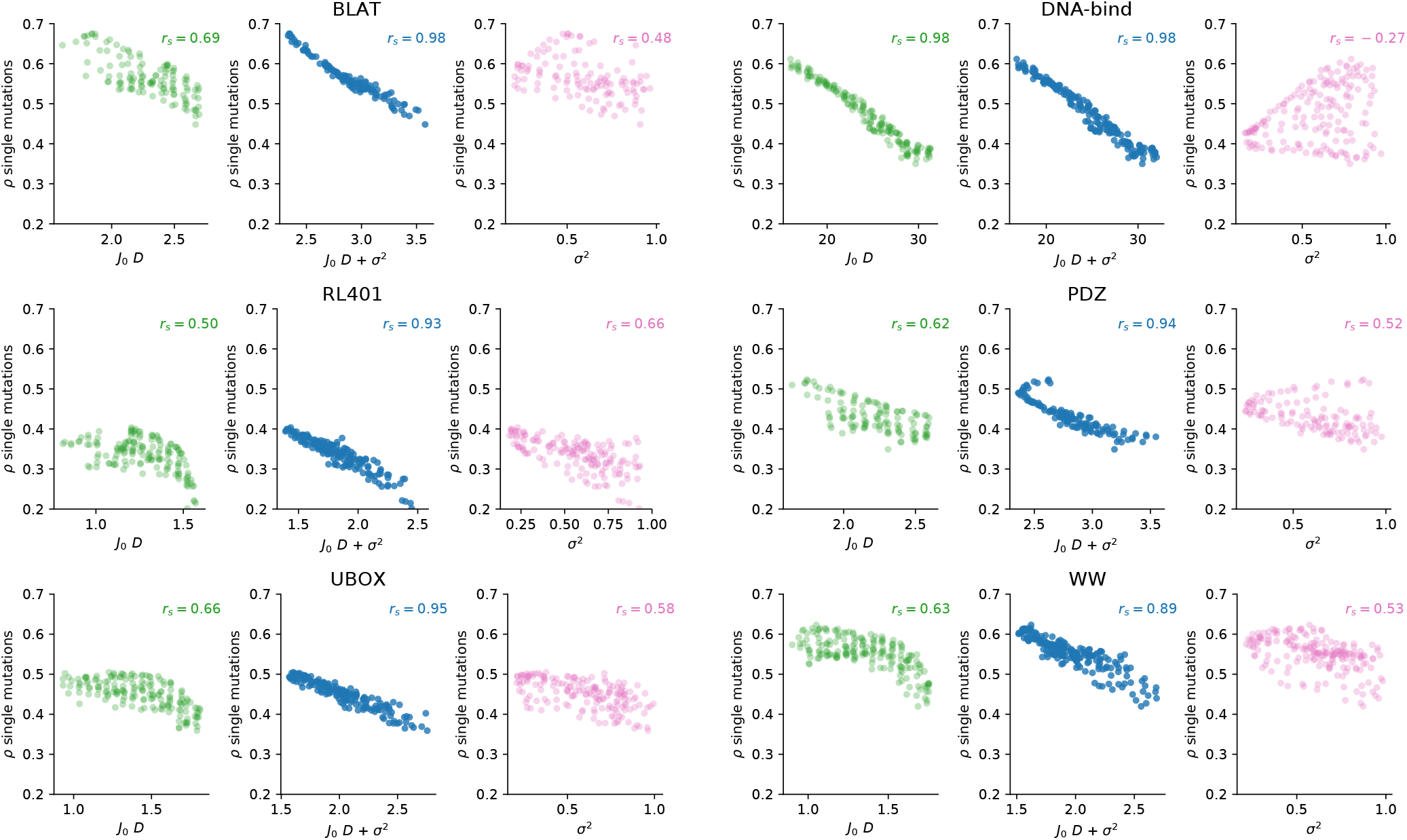
Same as Main text Fig. 4A&B for all protein families except RNA-Bind (shown in Main text Fig. 4A&B): predictive performance of single-point mutations using the independent-site models, as a function of the squared bias and variance estimated from the alignments, separately (left and right panels) and combined (central panel).

**Supplementary Figure 5.**
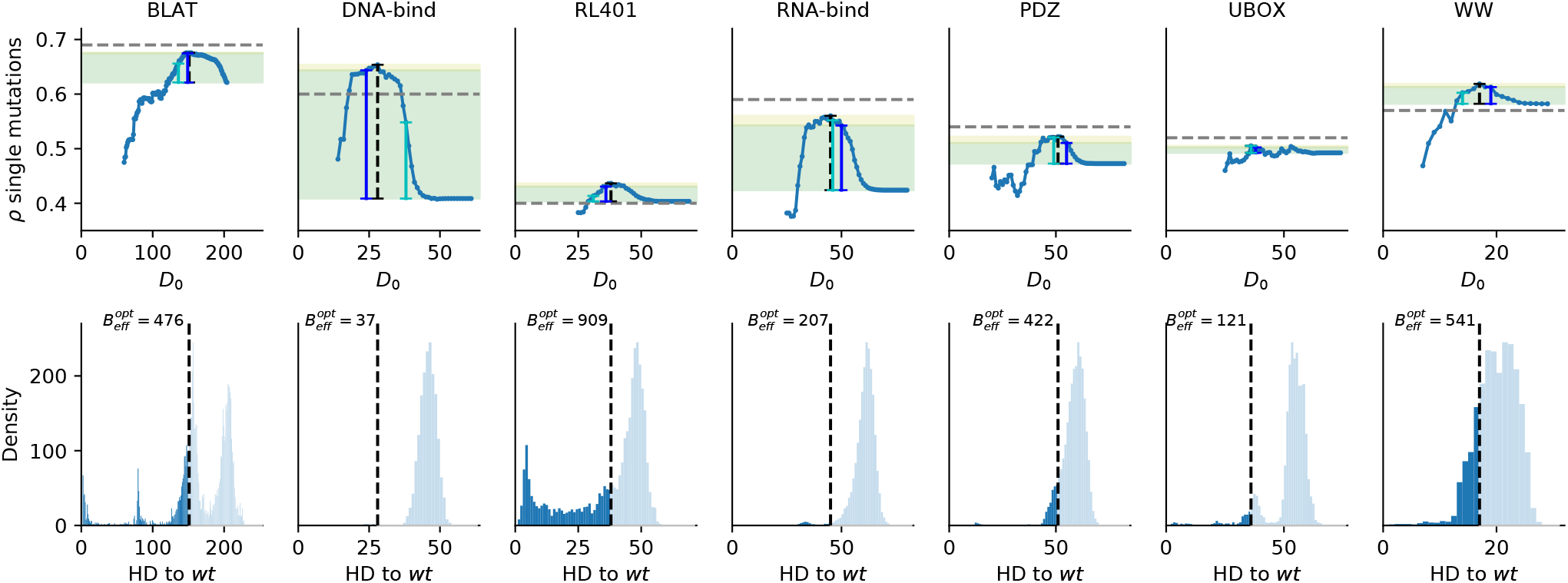
Top: single mutation prediction performance along the focusing axis (as a function of the cutoff distance *D*_0_) for the 7 studied protein families. Black dashed lines indicate the optimal cutoffs *d^opt^;* blue lines indicate the predicted cutoffs *d^bv^* by minimizing the linear sum of bias and variance; the light blue lines indicate the predicted cutoff from the signal-to-noise heuristic *d^snr^*. Green areas highlight the performance increase from the full alignment (*d_c_* = *N*) to the predicted cutoff *d^bv^*. Yellow areas indicate the remaining performance increase to the optimal cutoff *d^opt^*. Horizontal dashed grey lines indicate the performance reported in (37) with fully connected Potts model inferred by pseudo-likelihood. Bottom: distribution of the hamming distance to the wildtype *D* of sequences in the MSA. Black dashed lines indicate the optimal cutoff at which the best performance is reached. 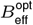 is the effective number of sequence remaining in the MSA at the optimal cutoff. Refer to Table 1 in Methods for the original number of sequences in the MSA.

**Supplementary Figure 6.**
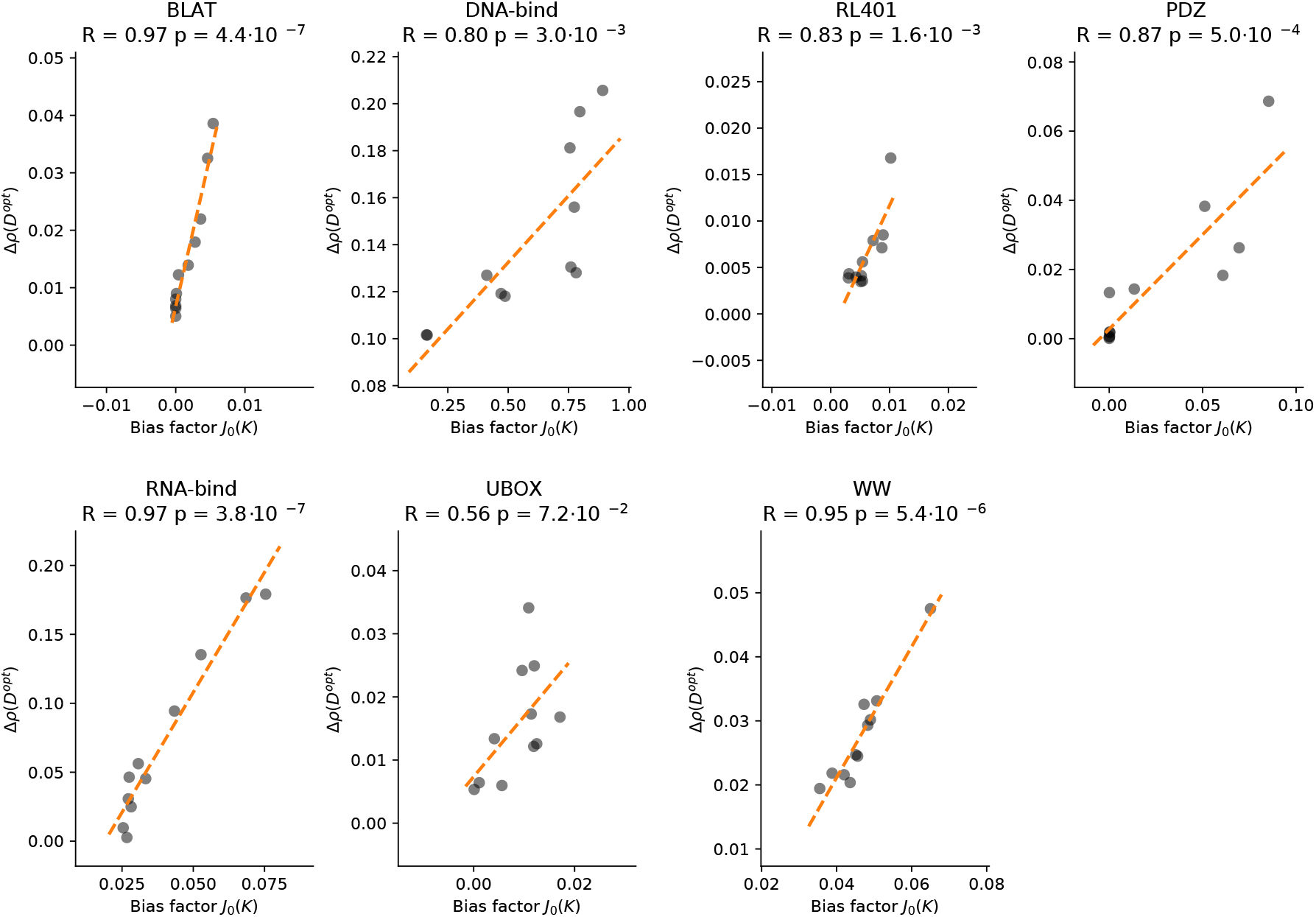
Relation between the bias factor *J*_0_(*K*) and improvement Δ*ρ*(*d^opt^*) for optimal focusing cutoff for the 7 studied protein families. For each family, *K* is varied between 0 and *N* (number of sites in the alignment).

## Footnotes

* Notice that the number of parameters to be inferred, *N_par_* = *N Q* + *K Q*^2^, where *Q* = 20 is the number of amino acids, grows quickly with *K* since *Q*^2^ = 400.

